# The minor spliceosome offers a therapeutically viable target for the treatment of a broad spectrum of cancers

**DOI:** 10.1101/2021.12.09.469623

**Authors:** Karen Doggett, Kimberly J Morgan, Stephen Mieruszynski, Benjamin B Williams, Anouk M Olthof, Alexandra L Garnham, Michael J G Milevskiy, Lachlan Whitehead, Janine Coates, Michael Buchert, Robert JJ O’Donoghue, Thomas E Hall, Zhiyuan Gong, Tracy L Putoczki, Matthias Ernst, Kate D Sutherland, Rahul N Kanadia, Joan K Heath

## Abstract

Minor splicing is a second splicing system required for the correct expression of ∼700 human minor intron-containing genes (MIGs). Many MIGs are expressed in vigorously proliferating cells and are frequently dysregulated in cancer including *BRAF, ERK, JNK* and *p38*. Minor splicing is carried out by the minor spliceosome which comprises several unique components, including a 65kDa protein encoded by *RNPC3*. We show that *Rnpc3* heterozygosity reduces tumour burden in a broad spectrum of *in vivo* cancer settings, without harming normal tissues. Using the collective power of zebrafish, mouse and human cancer models, we reveal a sequence of events connecting *Rnpc3* deficiency and impaired splicing of MIGs to DNA damage and activation of a Tp53-dependent transcriptional program that restricts tumour burden by inducing cell cycle arrest and apoptosis. Interrogation of human liver and lung cancer transcriptomes curated in TCGA revealed that the expression of many of the genes encoding protein components of the minor spliceosome is upregulated in these cancers. This is accompanied by upregulation of the expression of MIGs that are enriched in cell cycle and DNA damage pathways. These findings suggest that cancer cells can invoke mechanisms to increase the efficiency of minor splicing to support their high proliferation rates. Finally, Kaplan Meier survival analysis shows that highly expressed MIGs are frequently associated with poor patient survival. Taken together, these results indicate that the minor spliceosome offers a therapeutically viable target for the treatment of a broad spectrum of cancers.

## Introduction

Splicing is required to generate the correct RNA template for the synthesis of proteins and is an essential step in gene expression (1). For the most part, splicing is carried out by the major or U2-dependent spliceosome, which removes approximately 200,000 (99.65%) of introns. However, a second splicing system, known as the minor or U12-dependent spliceosome (2,3), is required to excise the remaining 0.35% of introns, which equates to 755 and 706 minor introns distributed across 699 human and 650 mouse genes, respectively (4). Minor introns are readily distinguished from major introns by the presence of two highly conserved sequence motifs: a 7bp sequence immediately downstream of the 5’ splice site (ss) and a branch point sequence (BPS) upstream of the 3’ss. They also lack the highly conserved polypyrimidine tract that is found upstream of the 3’ss of major introns (5). These distinctive properties of minor introns are recognized uniquely by the U11/U12 di-small nuclear ribonucleoprotein (di-snRNP) containing 2 small nuclear RNAs (snRNAs: U11, U12) and 8 proteins that are not utilized in the splicing of major introns (6-9). The next steps of minor splicing involve some additional unique components, namely, the U4ATAC and U6ATAC snRNAs and 4 recently described protein components (8). However, the downstream catalytic events that take place after intron recognition are mostly carried out by components shared with the major splicing machinery (7).

Despite their low frequency in the genome, the importance of minor introns is indicated by their evolutionary conservation in early eukaryotes (10), fungi, land plants and animals. Intriguingly, minor introns are not distributed randomly throughout the genome; instead, they are over-represented in sets of genes that perform DNA replication and RNA processing functions, including transcription, splicing, RNA quality control and translation (11). Of great interest in the context of cancer, minor introns are found in genes with established roles in tumourigenic pathways, including the proto-oncogenes, *BRAF* and *RAF1*, and 11 out of 14 mitogen-activated protein kinase (MAPK) family genes, including *ERK, JNK, p38* and their respective isoforms.

The direct consequence of impaired minor splicing is the accumulation of aberrant pre-mRNA transcripts exhibiting features such as minor intron retention and exon skipping (12). These events often generate frameshifts and premature stop codons in the coding sequence, leading to the production of truncated proteins or degradation of the improperly processed pre-mRNAs by nonsense mediated decay (NMD)(13). Severe consequences may result from the incorrect expression of minor intron containing genes (MIGs), including impaired cell growth, proliferation and survival, particularly during development (14-19).

Our interest in minor splicing was prompted by our discovery that *rnpc3* is required for the rapid growth and proliferation of cells in the developing digestive organs of larval zebrafish (20). *rnpc3* encodes a 65kDa RNA-binding protein (hereafter 65K) in the U11/U12 di-snRNP. We subsequently showed that *Rnpc3* is essential for pre-implantation murine development and that induced recombination of both alleles of the *Rnpc3* locus in adult mice severely impairs the homeostasis of the gastrointestinal epithelium, hematopoietic compartment and thymus (11). These results indicate a heightened requirement for *Rnpc3*/minor splicing in tissues undergoing rapid, continuous cell cycling, compared to quiescent tissues, and prompted us to think that minor splicing may be an Achilles’ heel of all highly proliferative tissues, including cancer. To test this hypothesis, we reduced *Rnpc3* expression in a broad spectrum of tumour-prone zebrafish and mouse cancer models and investigated the molecular consequences of knocking down *RNPC3* in human cancer cells.

## Results

### Heterozygous loss of *rnpc3* reduces tumour burden in a *kras*^*G12V*^-driven model of hepatocellular carcinoma

Zebrafish and mice carrying a single loss-of-function *rnpc3/Rnpc3* allele develop normally, achieve sexual maturity, and exhibit a normal lifespan (11,20). To test our hypothesis that *rnpc3*/65K is required for the growth and proliferation of cancer cells, we determined the impact of *rnpc3* heterozygosity in a zebrafish model of hepatocellular carcinoma (HCC) driven by a mutant *kras*^*G12V*^ transgene. We chose this model because the oncogenic properties of Kras^G12V^ reside in its ability to constitutively activate the Ras/Raf/Mek/Mapk pathway, several components of which are encoded by genes requiring minor splicing for their correct expression. In this *TO*(*kras*^*G12V*^) line, doxycycline induces the hepatocyte-specific expression of *EGFP*-*kras*^*G12V*^ causing hepatocyte hyperplasia and an increase in liver volume, which we quantified by two-photon microscopy (Figure S1) (21). As a control, we used the transgenic zebrafish line, *2-CLiP* (2-<u>C</u>olour <u>L</u>iver <u>P</u>ancreas) also known as *LiPan* (22), in which hepatocytes exhibit constitutive dsRed fluorescence but not the *EGFP*-*kras*^*G12V*^ transgene (Figure S1B).

Heterozygous *rnpc3* larvae contained 40% less *rnpc3* mRNA than *rnpc3*^*+/+*^ siblings at 7dpf (Figure 1A) but mean liver volume in *2-CLiP* larvae was unaffected by *rnpc3* genotype (Figure 1B, C). Dox-induced expression of *kras*^*G12V*^ in *rnpc3*^*+/+*^*;TO*(*kras*^*G12V*^) hepatocytes increased mean liver volume by >5-fold (Figure 1B, C, S1B). Remarkably, this increase was pared back to 3-fold in *TO(kras*^*G12V*^) larvae that were heterozygous for *rnpc3* (Figure 1B, C). Thus, heterozygous *rnpc3* is sufficient for normal liver growth during development but is rate-limiting for mutant *kras*^*G12V*^-driven hepatocyte hyperplasia. To determine if *rnpc3* heterozygosity affected cell cycle progression, we used EdU incorporation analysis to count the number of hepatocytes replicating DNA in S-phase. Compared to *TO(kras*^*G12V*^) larvae expressing WT *rnpc3*, the number of EdU-positive hepatocyte nuclei was reduced by 40% in *rnpc3*^*+/-*^*;TO(kras*^*G12V*^) larvae (Figure 1D, E). We then used zebrafish expressing an *annexinV-mkate* cell death reporter transgene to mark cells undergoing apoptosis (23). We observed a 50% increase in Annexin-mKate fluorescent foci in *TO(kras*^*G12V*^) livers on a heterozygous *rnpc3* background, compared to *TO(kras*^*G12V*^) livers expressing WT *rnpc3* (Figure 1F, G). These data indicate a vulnerability of *kras*^*G12V*^-expressing hepatocytes to *rnpc3* heterozygosity that reduces the number of cells in S-phase of the cell cycle and increases the frequency of cell death.

**Figure 1.**
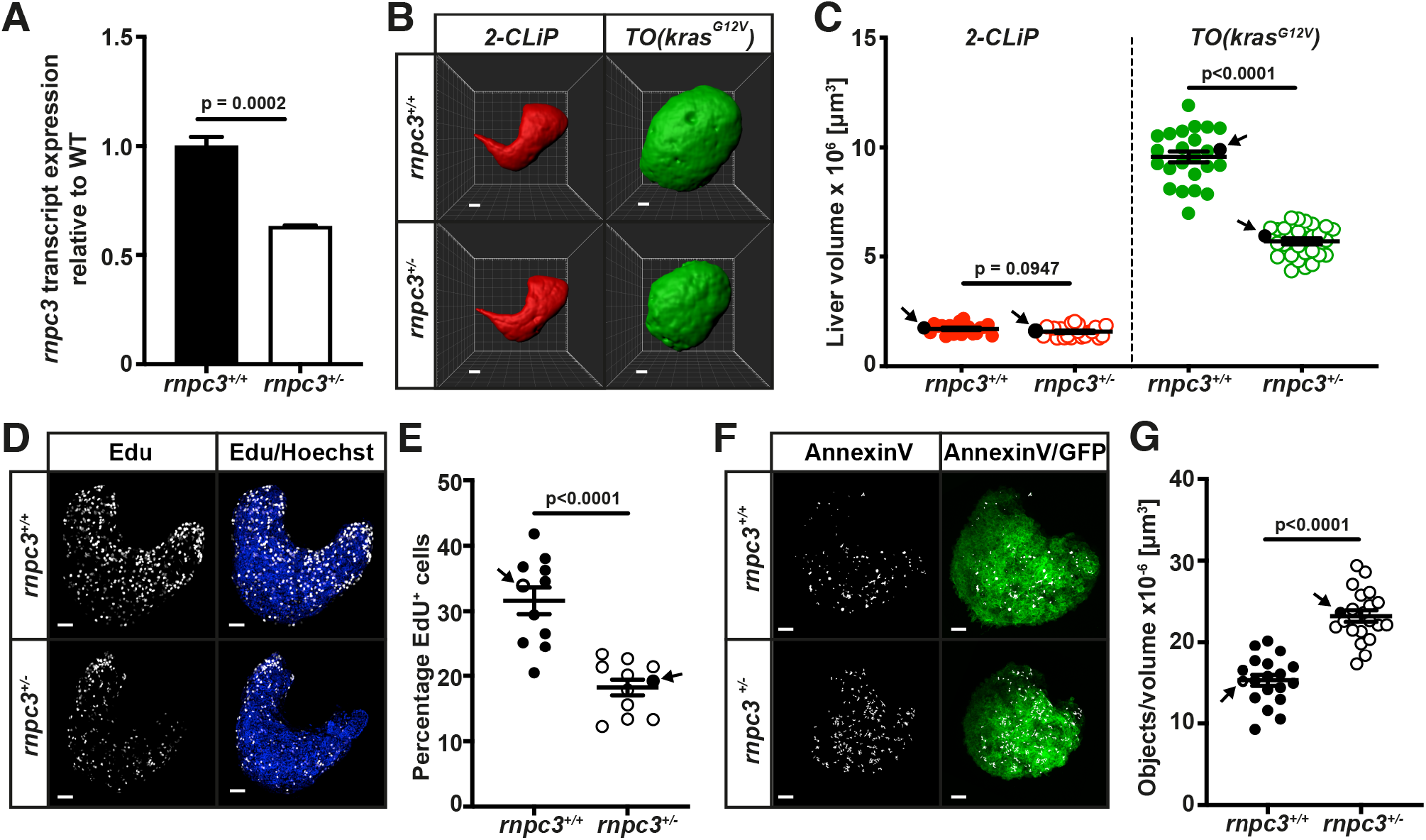
Heterozygous loss of *rnpc3* reduces tumour burden in a *kras*^*G12V*^*-*driven zebrafish model of hepatocellular carcinoma (HCC). **(A)** RT-qPCR analysis of *rnpc3* mRNA extracted from independent pools of *rnpc3*^*+/-*^ larvae compared to *rnpc3*^*+/+*^ larvae (n=3). **(B, C)** Mean liver volume in *2-CLiP* zebrafish at 7 days post-fertilization (dpf) is not affected by *rnpc3* genotype (red symbols, n=26 or n=29). Liver volume is increased approximately 5-fold in response to induced expression of the *TO(kras*^*G12V*^) transgene (solid green circles, n=24), compared to normal livers on the *2-CLiP* background (solid red circles, n=28). Heterozygous loss of *rnpc3* (open green circles) significantly restricts the *TO(kras*^*G12V*^*)-*driven increase in liver volume. Black arrows on the graphs indicate the data points (black symbols) that correspond to the representative images shown in **(B). (D, E)** The abundance of Hoechst 33342-positive hepatocyte nuclei that incorporated EdU (white dots) in S phase of the cell cycle is reduced significantly in *TO(kras*^*G12V*^) livers on a heterozygous *rnpc3* background, compared to *TO(kras*^*G12V*^) livers on a WT *rnpc3* background (n=11). **(F, G)** Foci of fluorescence show AnnexinV-mKate molecules bound to apoptotic cells (white objects). These are more abundant in *TO(kras*^*G12V*^) livers on a heterozygous *rnpc3* background (n=20), compared to *TO(kras*^*G12V*^) livers on a WT *rnpc3* background (n=19). Data are represented as mean ± SEM. Significance was assessed using a Student’s *t* test, P<0.05. Scale bar in **A** is 25 µm and 50 μm in **D** and **F**.

### Heterozygous loss of *Rnpc3* reduces tumour burden in a *Kras*^*G12D*^-driven mouse model of lung adenocarcinoma

Next, we determined whether *Rnpc3* heterozygosity retards the growth of mutant *Kras*-driven cancer in a well-characterized mouse model of lung adenocarcinoma. *Kras*^*LSLG12D/+*^ mice contain one WT *Kras* allele and one inducible (*loxP*-STOP-*loxP*) *Kras*^*G12D/+*^ allele knocked-in to the *Kras* locus (24). In the absence of Cre-mediated recombination, mice are heterozygous for WT *Kras*. We induced expression of oncogenic *Kras*^*G12D*^ in lung epithelial cells by intranasal delivery of adenoviral Cre recombinase (AdCre) (25). Histological examination of the lungs of *Rpnc3*^+/+^;*Kras*^*G12D*^ mice, 180 d after induction of oncogenic *Kras*^*G12D*^ expression, revealed the presence of multifocal preneoplastic epithelial lesions, classified as either atypical adenomatous hyperplasia (AAH, arrows), or more advanced papillary adenomas and micro-adenocarcinomas (arrowhead; Figure 2A and 2B), reminiscent of early to intermediate stages of the human disease. These lesions stained robustly for pERK1/2 (Figure 2C) and the *Rnpc3*-encoded protein, 65K (Figure 2D). In contrast, the lungs from *Rpnc3*^*+/-*^;*Kras*^*G12D*^ mice exhibited smaller lesions than lungs from *Rnpc3*^*+/+*^;*Kras*^*G12D*^ mice (compare Figure 2A and 2E). Histopathological analysis confirmed that most of these lesions were AAH (arrows, Figure 2E, F) with few papillary adenomas/adenocarcinomas. The smaller lesions also expressed pERK (Figure 2G) and 65K (Figure 2H). Quantification of tumour burden revealed a significant decrease in the percentage of total lung area occupied by hyperplastic lesions in heterozygous *Rnpc3* mice compared to WT *Rnpc3* mice (Figure 2I) and a reduction in the number of lesions per mm^2^ of lung area (Figure 2J).

**Figure 2.**
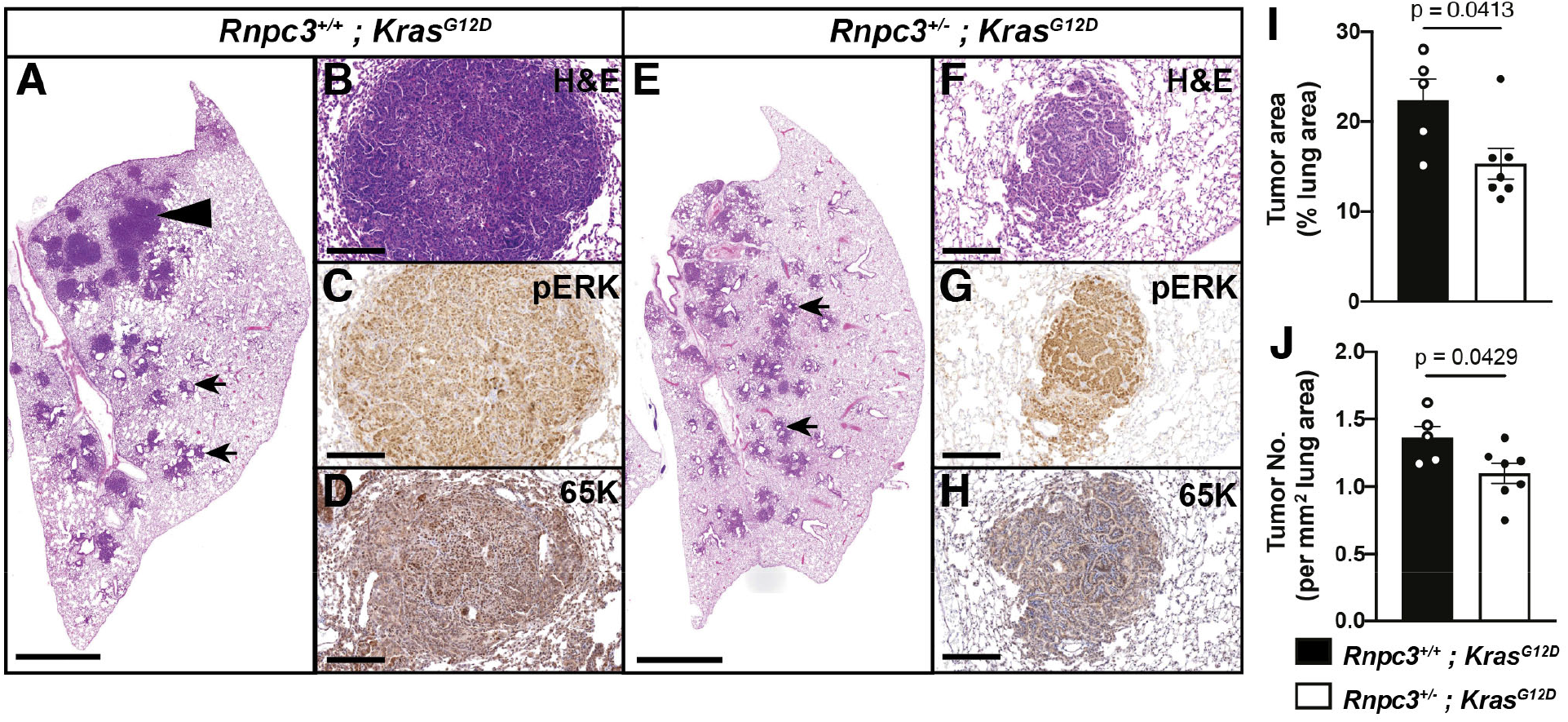
*Rnpc3* heterozygosity reduces tumour burden in a mouse model of lung adenocarcinoma. *Kras*^*G12D*^ expression was induced in the lung airway epithelium of mice by intranasal administration of adenoviral Cre recombinase (AdCre). Histological examination of the lungs was performed 180 d later. **(A, B)** Representative hematoxylin and eosin-stained lung sections from *Rpnc3*^*+/+*^*;Kras*^*G12D*^ mice. Arrows and arrowhead indicate foci of atypical adenomatous hyperplasia (AAH) and adenoma, respectively. These lesions are pERK **(C)** and 65K **(D)** positive. **(E, F)** In *Rpnc3*^*+/-*^*;Kras*^*G12D*^ mice the hyperplastic lesions are smaller (arrows) and are also pERK **(G)** and 65K positive **(H). (I)** The area of hyperplasia in sections of lung, expressed as a percentage of total lung area, was significantly decreased in *Rnpc3*^*+/-*^ mice compared to *Rnpc3*^*+/+*^ mice. **(J)** The number of hyperplastic lesions per mm^2^ lung area. Scale bar in **A** and **E** is 2 mm, scale bar in **B**-**D** and **F**-**H** is 200 μm. Results are expressed as mean ± SEM, n=5 or 7 per genotype. Significance was assessed using a Student’s *t* test, P<0.05.

We also crossed the same lung cancer model onto a *Rnpc3*^*lox/lox*^ background (11). These mice differ from the previous mice because they are not constitutively heterozygous for *Rnpc3*. Thus, when we administered AdCre intranasally, we recombined the conditional *Rnpc3*^*lox/lox*^ and *Kras*^*G12D*^ alleles simultaneously in the same cells. Histological examination of the lungs 90 d after AdCre delivery revealed a significant reduction in the tumour area and the number of AAH lesions in *Rnpc3*^*lox/lox*^*;Kras*^*G12D*^ lungs, compared to *Rnpc3*^*+/+*^*;Kras*^*G12D*^ control lungs (Figure S2A-D), demonstrating that the impact of reducing *Rnpc3* expression during lung tumourigenesis is intrinsic to cancer cells expressing *Kras*^*G12D*^.

### Heterozygous loss of *Rnpc3* reduces tumour burden in a STAT3-driven model of gastric cancer

To explore whether heterozygous *Rnpc3* reduces tumour burden in cancers not driven by *Kras* oncogenes, we used a mouse model of gastric cancer in which adenomas develop due to a mutation in GP130, the common signal transducer of the IL6 family of cytokines. In this *Gp130*^*F/F*^ model, mice harbor a knock-in Y→F missense mutation at codon 757 in both alleles of *Gp130* (*Gp130*^*Y757F/Y757F*^). This disrupts a SOCS3-mediated negative feedback mechanism leading to persistent activation of STAT3 and the spontaneous, 100% penetrant, development of adenomas in the glandular epithelium of the stomach by 100 d of age (26,27). Mice with a *Rnpc3*^*+/-*^;*Gp130*^*F/F*^ genotype exhibited a marked reduction in total adenoma weight compared to *Rnpc3*^*+/+*^;*Gp130*^*F/F*^ mice (Figure 3A-C) at 100 d, with a more marked reduction in the weight of adenomas harvested from the proximal glandular epithelium (corpus; arrowheads) compared to the distal glandular epithelium (antrum; arrows). Indeed, approximately 40% of *Rnpc3*^*+/-*^;G*p130*^*F/F*^ mice displayed a complete absence of corpus adenomas at 100 d (compare 3A, G with 3B, H) and this observation was sustained when the mice were aged to 180 d (Figure 3D-F). We observed pERK1/2 staining at the luminal edge of adenomas in both the corpus and antral regions (Figure 3I, J), indicating that MAPK signalling was active at the proliferative front of the growing adenomas. Using Western blot analysis, we quantitated the abundance of pERK1/2 proteins in *Rnpc3*^*+/-*^ and *Rnpc3*^*+/+*^ adenomas and observed a 35% reduction in the pERK1/2 signals in *Rnpc3*^*+/-*^ cells (Figure 3K, L), indicating attenuated activation of the MAPK pathway.

**Figure 3.**
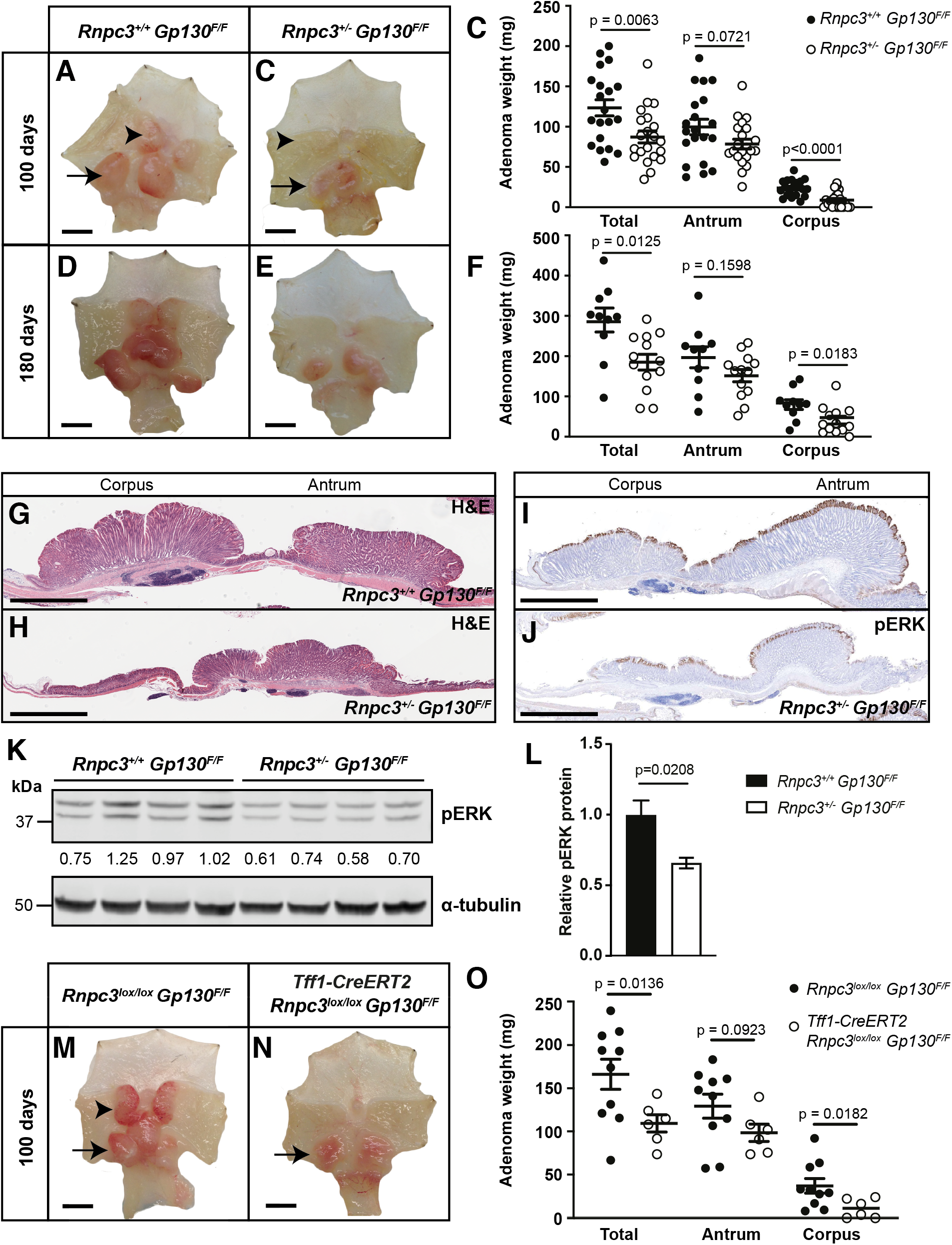
Heterozygous loss and induced deletion of *Rnpc3* reduces adenoma burden in a STAT3-driven model of gastric cancer. **(A)** Adenomas in the corpus (arrowhead) and antral (arrow) regions of the glandular stomach of *Rnpc3*^*+/+*^*;Gp130*^*F/F*^ and **(B)** *Rnpc3*^*+/-*^*;Gp130*^*F/F*^ mice at 100 d of age. **(C)** Total adenoma weight was significantly reduced in the stomachs of *Rnpc3*^*+/-*^mice (white circles, n=21) compared to *Rnpc3*^*+/+*^ mice (black circles, n=20), most notably in the corpus region. **(D-F)** Representative images of adenomas harvested from the gastric mucosa of *Rnpc3*^*+/-*^*;Gp130*^*F/F*^ mice at 180 d indicate a reduction in tumour burden, again with a more marked impact on the corpus region **(F)**. Data are expressed as mean ± SEM, n=10 or 14 per genotype. Significance was assessed with a Student’s *t* test with Welch’s correction. **(G, H)** Histological sections of the glandular stomach stained with hematoxylin and eosin reveal a reduction in adenoma burden in *Rnpc3*^*+/-*^ mice at 100d. **(I, J)** Immunocytochemical localization of pERK1/2 indicates active MAPK signaling at the luminal surface of adenomas in 100d old *Gp130*^*F/F*^ mice. **(K)** Western blot analysis of pERK1/2 proteins in individual antral adenomas collected from 4 individual *Rnpc3*^*+/+*^*;Gp130*^*F/F*^ and *Rnpc3*^*+/-*^*;Gp130*^*F/F*^ mice at 100d. Values shown are normalized by reference to the α-tubulin loading control and relative to *Rnpc3*^*+/+*^*;Gp130*^*F/F*^ samples. **(L)** Quantification of pERK1/2 protein abundance shown in **K**. Data are expressed as mean ± SEM, n=4. Significance was assessed with a Student’s *t* test. **(M)** Representative stomach from TMX-treated *Rnpc3*^*lox/lox*^*;Gp130*^*F/F*^ mice (no *Tff1-CreERT2* transgene) shows adenomas in the corpus (arrowhead) and antrum (arrow). **(N)** Representative stomach from TMX-treated *Rnpc3*^*lox/lox*^*;Gp130*^*F/F*^ mice harbouring the *Tff1-CreERT2* transgene shows conspicuously less adenomatous tissue. **(O)** Quantitation of adenoma weight in TMX-treated *Rnpc3*^*lox/lox*^*;Gp130*^*F/F*^ mice in the presence and absence of the *Tff1-CreERT2* transgene. Data are expressed as mean ± SEM, n=6 or 10 per genotype. Significance was assessed using an unpaired Student’s *t* test with Welch’s correction. Scale bar in **A, B, D, E, M** and **N** is 5 mm. Scale bar in **G-J** is 2 mm.

To determine whether inducing loss of *Rnpc3* expression in already established gastric adenomas still impacted on tumour growth (compared to constitutive heterozygosity), we crossed mice carrying conditional *Rnpc3*^*lox*^ alleles with *Gp130*^*F/F*^ mice also carrying a *Trefoil factor 1 (Tff1)-CreERT2* BAC transgene that confers tamoxifen (TMX)-inducible, glandular stomach-specific, Cre recombinase activity (28). We recombined *Rnpc3*^*lox*^ alleles specifically in the gastric epithelium by oral gavage of TMX to mice on days 56 and 57, well after adenoma formation has initiated (29) (Figure S3A). Recombination of *Rnpc3*^*lox*^ alleles produced a reduction in total adenoma burden, again with a more marked effect on the corpus region than the antrum, compared to TMX-treated mice with no *Tff1-CreERT2* BAC transgene (Figure 3M-O).

Antral adenomas harvested from TMX-treated *Tff1-CreERT2;Rnpc3*^*lox/lox*^*;Gp130*^*F/F*^ mice contained 50% fewer *Rnpc3* transcripts compared to antral adenomas from TMX-treated *Rnpc3*^*lox/lox*^*;Gp130*^*F/F*^ (no *Cre*-transgene) controls (Figure S3B). Individual adenomas from TMX-treated *Tff1-CreERT2;Rnpc3*^*lox/lox*^*;Gp130*^*F/F*^ mice contained cells with either a *Rnpc3*^*lox/lox*^ or *Rnpc3*^*lox/Δ*^ genotype, where *Δ* represents a deleted (null) allele (Figure S3C), corresponding to cells that evaded recombination of one or both *Rnpc3*^*lox*^ alleles. No cells exhibited a *Rnpc3*^*Δ/Δ*^ (completely null) genotype, suggesting that *Rnpc3*^*Δ/Δ*^ cells died. We interpret the reduction in adenoma burden observed in TMX-treated *Tff1-CreERT2;Rnpc3*^*lox/lox*^*;Gp130*^*F/F*^ mice to be due to the combined effect of two induced genotypes: the *Rnpc3*^*lox/Δ*^ (heterozygous) genotype causing reduced proliferation and the *Rnpc3*^*Δ/Δ*^ (homozygous null) genotype causing cell death.

### Disruption of the *Rnpc3* locus in AML cells causes impaired minor splicing and prolonged survival of mice

Our findings from three different solid tumour models suggest that hyperproliferative cancers require efficient minor splicing to proliferate and survive. To test this further, we employed a tractable model of blood cancer. Translocations of the Mixed Lineage Leukemia (*MLL*) gene, encoding a histone methyl transferase, are associated with a subset of aggressive AMLs with poor prognosis (30). The translocation t(11;19) fuses the genes *MLL* and *ENL* to encode the leukemogenic oncoprotein MLL-ENL (31), a recurrent feature of a subgroup of acute leukemias occurring in infants. We used cells containing a retroviral construct encoding the human MLL-ENL oncogene fused to GFP to model AML in mice (32) (Figure 4A). Foetal liver cells harvested from mice harbouring conditional *Rnpc3*^*lox/lox*^ alleles and a *UBC-CreERT2* transgene (11) were transformed with retroviral MLL-ENL-GFP to generate a population of cells that was used to reconstitute the hematopoietic system of irradiated WT mice (Figure 4A). As each mouse reached the ethical endpoint (latency = 42-81 d), AML cells were harvested from the bone marrow and spleen and transplanted into unirradiated WT hosts where they generated a more aggressive AML with a shorter latency (17-55 d; Figure 4A). These AML cells were harvested from the bone marrow and were either treated with 4-hydroxy tamoxifen (4-OHT) *in vitro* to assess the impact of recombining the *Rnpc3*^*lox/lox*^ locus on the efficiency of minor splicing and cell viability in culture, or transplanted for the third time into recipient WT mice and treated with TMX 13 and 14 d later to determine the impact of recombining the *Rnpc3*^*lox/lox*^ locus on aggressively dividing AML cells *in vivo*.

**Figure 4.**
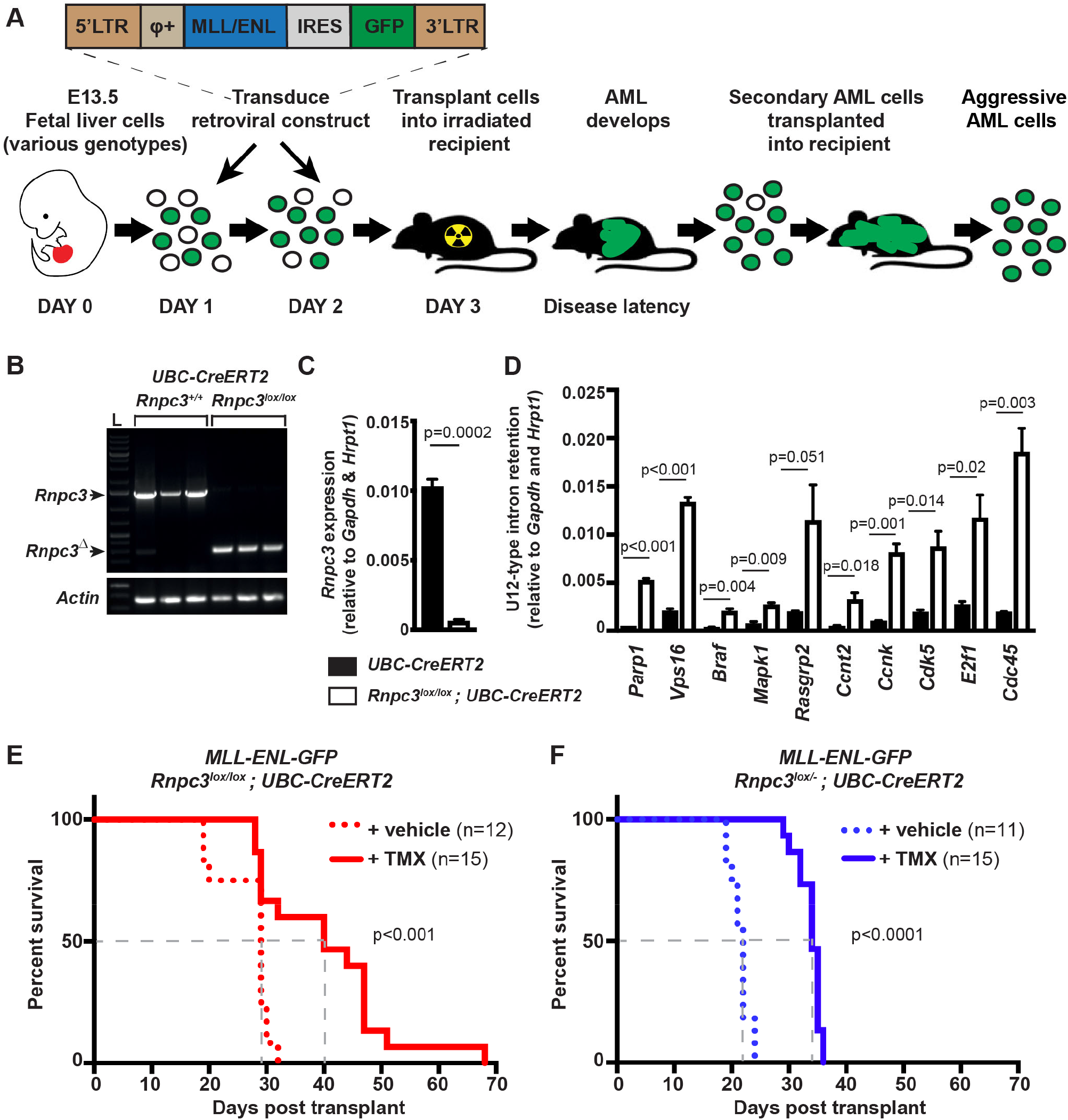
Disruption of the *Rnpc3* locus in AML cells causes impaired minor splicing and prolonged survival of mice. **(A)** Schematic diagram depicting the generation of AML cell lines from pooled foetal liver cells harvested from E13.5 mice of various genotypes (either WT, *UBC-CreERT2, Rnpc3*^*lox/lox*^*;UBC-CreERT2* or *Rnpc3*^*lox/-*^*;UBC-CreERT2*) and transduced on successive days with a retroviral construct encoding the MLL-ENL fusion protein and GFP. Primary *MLL-ENL-*transduced cells were transplanted intravenously into irradiated WT mice and propagated until the ethical endpoint of the experiment. *MLL-ENL* AML cells were harvested from the bone marrow and transplanted into (unirradiated) WT recipient mice for further propagation, enriching the proportion of MLL-ENL AML cells in the bone marrow and other hematopoietic organs. These secondary MLL-ENL AML cell lines were then either cultured *in vitro* for 72 h in medium containing 4-OHT (**B-D**) or transplanted again into recipient mice (**E, F**). **(B)** RT-PCR shows that 4-OHT-treatment of *Rnpc3*^*lox/lox*^;*UBC-CreERT2* AML cells *in vitro* achieves essentially 100% recombination of *loxP*-flanked *Rnpc3* alleles. L=Ladder. **(C)** RT-qPCR reveals a >10-fold reduction in *Rnpc3* transcripts in 4-OHT-treated *Rnpc3*^*lox/lox*^;*UBC-CreERT2* cells compared to 4-OHT-treated *UBC-CreERT2* cells. **(D)** Reduced expression of *Rnpc3* causes a significant impairment in minor splicing efficiency, resulting in the detection of amplicons containing retained minor introns. Results are expressed as the mean of 3 independent secondary AML cell lines per genotype ± SEM. Significance was assessed using a two-tailed, unpaired Students’ *t* test P<0.05. **(E)** Kaplan-Meier plot of WT mice harbouring tertiary transplants of *Rnpc3*^*lox/lox*^;*UBC-CreERT2* AML cells and treated with TMX or vehicle 13 and 14 days later to induce recombination of the *Rnpc3* locus. Mice harbouring *Rnpc3*^*lox/lox*^;*UBC-CreERT2* AML cells treated with TMX survived significantly longer (red line, median survival 40 d, n=15) than vehicle-treated mice transplanted with the same cells (red dotted line, median survival 29 d, n=12). **(F)** Mice bearing tertiary transplants of *Rnpc3*^*lox/-*^;*UBC-CreERT2* AML cells treated with TMX survived significantly longer (blue line, median survival 34 d, n=15) than vehicle-treated mice harbouring the same tertiary AML cells (blue dotted line, median survival 22 d, n=11). Significance was assessed with a Mantel-Cox test.

*Rnpc3*^*lox*^ alleles in AML cells carrying the *UBC-CreERT2* transgene underwent almost complete recombination after treatment with 4-OHT *in vitro* (Figure 4B). This resulted in a 95% loss of *Rnpc3* mRNA expression at 72 h (Figure 4C) and a marked increase in cell death at 96 and 120 h, as shown by flow cytometry analysis of AnnexinV/Propidium iodine staining (Figure S4A, expressed graphically in S4B). The impact of loss of *Rnpc3* expression on the efficiency of minor splicing in secondary AML cells was assessed by RT-qPCR using primers designed to amplify retained minor introns. We analysed transcripts relevant to DNA repair (*Parp1*), MAPK signalling (*Braf, Mapk1, Rasgrp2*) and cell cycle progression (*Ccnt2, Ccnk, Cdk5, E2f1, Cdc45*) and *Vps16*. In 9 out of 10 cases, transcripts from AML cells with a disrupted *Rnpc3* locus exhibited significantly elevated minor intron retention compared to cells in which the *Rnpc3* locus remained intact (Figure 4D), showing that disrupting *Rnpc3* expression in AML cells impairs the efficiency of minor splicing and leads to cell death.

In an approach akin to a targeted therapeutic intervention, we recombined the *Rnpc3* locus *in vivo*. In the absence of TMX treatment, mice harbouring *Rnpc3*^*lox/lox*^*;UBC-CreERT2* AML cells exhibit a median survival of 29 d. However, disrupting the *Rnpc3* locus at 13 and 14 d with TMX extended the median survival to 40 d (38% increase; Figure 4E; Figure S4C for additional controls). Likewise, when WT recipient mice were transplanted with *Rnpc3*^*lox/-*^;*UBC-CreERT2* AML cells and treated with TMX, the median survival of the mice was extended by >50% (34 d compared to 22 d; Figure 4F). Ultimately, all mice developed splenomegaly (Figure S4D) and succumbed to AML with an immature myeloid (Mac-1^+^/Gr-1^-^) phenotype (Figure S4E). Genomic analysis of the TMX-treated tertiary transplanted AML cells revealed the presence of both unrecombined (*Rnpc3*^*lox*^) and deleted (*Rnpc3*^*Δ*^) alleles in AML cells derived from *Rnpc3*^*lox/lox*^ mice, indicating that TMX treatment did not achieve complete recombination of the *Rnpc3* locus *in vivo* (Figure S4F). Only the unrecombined *Rnpc3*^*lox*^ allele was detected in AML cells derived from *Rnpc3*^*lox/-*^;*UBC-CreERT2* mice, suggesting that AML cells that acquired a *Rnpc3*^*Δ/−*^ genotype died.

### *rnpc3* heterozygosity combines with *kras*^*G12V*^ to activate a Tp53 DNA damage response that restricts tumour burden

In our zebrafish model of *kras*^*G12V*^-driven HCC, *rnpc3* heterozygosity reduced tumour burden by decreasing the number of cells in S-phase of the cell cycle and increasing cell death (Figure 1). Since these responses are reminiscent of activation of the tumour suppressor functions of Tp53, we measured the levels of Tp53 protein in lysates of micro-dissected livers expressing the *kras*^*G12V*^ transgene or not. No Tp53 signal was obtained from non-*kras*^*G12V*^-expressing livers, or larval remains after liver removal, irrespective of *rnpc3* genotype. We detected a weak Tp53 signal in extracts of *kras*^*G12V*^-expressing livers on a WT *rnpc3* background, and a markedly stronger Tp53 signal (>2.4-fold) in *kras*^*G12V*^-expressing livers from heterozygous *rnpc3* larvae (Figure 5A). To test whether Tp53 was responsible for the reduction in tumour burden in *rnpc3*^*+/-*^;*kras*^*G12V*^ livers, we repeated the experiment in the presence and absence of zebrafish *tp53*^*M214K*^ alleles (hereafter *tp53*^*m*^), which encode missense mutations in the DNA-binding domain of Tp53 that abrogate Tp53 function (33). Mean liver volume in *rnpc3*^*+/+*^*;TO(kras*^*G12V*^)^*T/+*^ larvae on a *tp53*^*m/m*^ background was increased (+33%) compared to larvae with WT *tp53* expression (Figure 5B, C), indicating that Tp53 function restrains tumour growth in this model. This was more evident in *rnpc3*^*+/-*^*;TO(kras*^*G12V*^)^*T/+*^ larvae, where liver volume increased by >2-fold in mutant *tp53*^*m/m*^ larvae compared to WT *tp53* larvae. Indeed, in the absence of functional Tp53, *rnpc3* heterozygosity reduced liver volume by only 8% compared to larvae with WT *rnpc3*, showing that the capacity of heterozygous *rnpc3* to restrain tumour growth relies heavily on activating Tp53 function.

**Figure 5.**
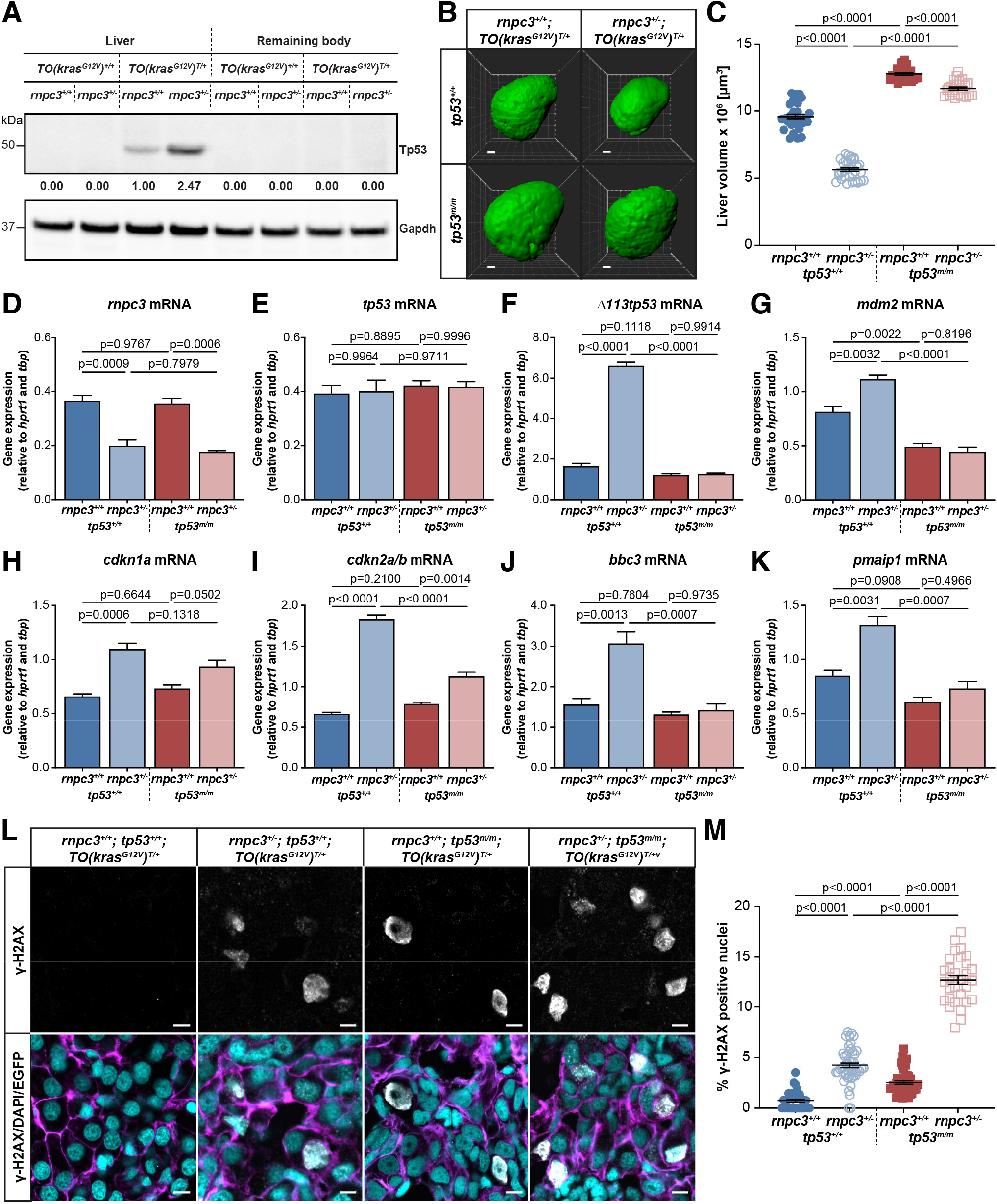
*rnpc3* heterozygosity combines with *kras*^*G12V*^ to activate a Tp53 DNA damage response that restricts tumour burden. **(A)** Western blot showing Tp53 protein signals in lysates of *TO(kras*^*G12V*^) larvae of the indicated *rnpc3* genotype. Values shown are normalized by reference to the Gapdh loading control and compared with the Tp53 signal in lane 3, which was set at 1. **(B)** Representative three-dimensional reconstructions of *TO(kras*^*G12V*^)^*T/+*^ livers of the indicated *rnpc3* and *tp53* genotypes. Scale bar is 25 µm. **(C)** Impact of *rnpc3* heterozygosity and homozygous *tp53* mutation on liver volume in *TO(kras*^*G12V*^)^*T/+*^ larvae. Error bars represent mean ± SEM, n=28 for *rnpc3*^*+/+*^; *tp53*^*+/+*^, n=32 for *rnpc3*^*+/-*^; *tp53*^*+/+*^, n=27 for *rnpc3*^*+/+*^; *tp53*^*m/m*^, n=24 for *rnpc3*^*+/-*^; *tp53*^*m/m*^. Scale bar is 25µm. **(D-K)** RT-qPCR analysis of gene expression in *TO(kras*^*G12V*^)^*T/+*^ dissected livers of the indicated *rnpc3* and *tp53* genotypes. Data are expressed as mean ± SEM, n=3. **(L)** Representative Airyscan imaging of liver cryosections of liver from *TO(kras*^*G12V*^)^*T/+*^ larvae of the indicated *rnpc3* and *tp53* genotype stained with γ-H2AX antibody (white) marking DNA double-strand breaks, DAPI (cyan) marking DNA, and EGFP-Kras^G12V^ (magenta) marking the cell membrane. Scale bar is 5µm. **(M)** Quantification of the percentage of hepatocytes positive for γ-H2AX. Data are expressed as mean ± SEM, n=32 for *rnpc3*^*+/+*^; *tp53*^*+/+*^, n=47, for *rnpc3*^*+/-*^; *tp53*^*+/+*^ n=46 for *rnpc3*^*+/+*^; *tp53*^*m/m*^, n=30 for *rnpc3*^*+/-*^; *tp53*^*m/m*^. Significance was tested using a one-way ANOVA with Tukey’s multiple comparisons test.

We used RT-qPCR to determine expression of candidate genes in *kras*^*G12V*^-expressing livers in the presence and absence of Tp53 function. *tp53* mRNA expression was not altered by *rnpc3* genotype (Figure 5E), indicating that post-transcriptional mechanisms were responsible for the different levels of Tp53 protein we detected between larvae that were WT or heterozygous for *rnpc3* (Figure 5A). That Tp53 was transcriptionally active was shown by the upregulated mRNA expression of canonical Tp53 target genes, Δ*113tp53* and *mdm2* (4.0-fold and 1.4-fold), respectively (Figure 5F, G). We then measured the expression of two cell cycle arrest genes, *cdkn1a* and *cdkn2a/b* (encoding p21 and p14^ARF^/p16^INK4A^, respectively) and two Bcl2 family genes that promote mitochondrial apoptosis, *pmaip1* and *bbc3* (Figure 5H-K). All four genes were significantly upregulated in *rnpc3*^*+/-*^*;tp53*^*+/+*^*;TO(kras*^*G12V*^)^*T/+*^ livers compared to *rnpc3*^*+/+*^*;tp53*^*+/+*^*;TO(kras*^*G12V*^)^*T/+*^ livers, consistent with Tp53 playing a role in reducing tumour burden in heterozygous *rnpc3* livers.

Since DNA damage is a strong signal for Tp53 activation, we used antibodies to γ-H2AX to look for DNA double strand (ds) breaks in DAPI-stained cryosections of liver (Figure 5L). We found very few hepatocyte nuclei contained DNA ds breaks (0.8%) in *TO(kras*^*G12V*^)^*T/+*^ larvae that were WT for *rnpc3* and *tp53* (Figure 5M). However, the frequency of DNA ds breaks increased >3-fold in the absence of Tp53 function. In heterozygous *rnpc3* hepatocytes, the percentage of γ-H2AX positive nuclei increased 5-fold, compared to WT *rnpc3* hepatocytes, showing that the combination of *kras*^*G12V*^ and heterozygous *rnpc3* expression increased DNA damage. Most striking, the percentage of γ-H2AX positive hepatocytes increased a further 3-fold (to 12.7%) in heterozygous *rnpc3* hepatocytes in the absence of Tp53 function, demonstrating that *rnpc3*^*+/-*^*;kras*^*G12V*^ hyperplastic hepatocytes incur considerable DNA damage, which remains unrectified in the absence of functional Tp53 (Figure 1D-G, Figure 5H-K).

### Knockdown of *RNPC3* reduces growth of human cancer cells

We chose A549 cells, derived from a lung adenocarcinoma, and HeLa cells to investigate the impact of *Rnpc3*-depletion on the growth of human cancer cells *in vitro*. A549 cells carry a *KRAS*^*G12S*^ mutation, akin to the *kras*^*G12V*^ and *KRAS*^*G12D*^ mutations in the *in vivo* mutant *Kras* cancer models we showed are sensitive to heterozygous *Rnpc3*. We transfected these cells with siRNAs targeting *RNPC3* and *PDCD7* to achieve levels of depletion (between 50-100%) that are more relevant to therapeutic intervention than can be achieved genetically with animal models. *PDCD7* encodes U11/U12 59K, another unique protein component of the minor spliceosome. Together with 48K and 65K, 59K forms a ‘molecular bridge’ between the conserved 5’ss and BPS sequences of minor introns prior to splicing (34). We used non-targeted (NT) siRNA for controls.

Following transfection with siRNAs, we tracked confluency of A549 and HeLa cells over time in an IncuCyte Live-Cell analysis system. Compared to siNT-treated cells, A549 cells transfected with *RNPC3s* and *PDCD7* siRNAs grew more slowly from 24 h onwards. By 72 h cell confluency was reduced by 35.8-37.2% and 31.1-36.8% in response to *RNPC3* and *PDCD7* knockdown, respectively (Figure 6A, B; Figure S5A, B). At 72 h, si*RNPC3* reduced *RNPC3* mRNA expression in A549 cells by >70% (Figure 6C) and reduced the 65K protein signal by 45-62% (Figure S6B, C), while si*PDCD7* produced a >85% reduction in *PDCD7* mRNA expression (Figure S5C). HeLa cells responded to *RNPC3* and *PDCD7* knockdown in a comparable fashion to A549 cells after a 48 h incubation (Figure S5E-H). Both cell lines increased *RNPC3* expression in response to *PDCD7* knockdown (Figure S5G), likely attempting to compensate for *PDCD7* deficiency.

**Figure 6.**
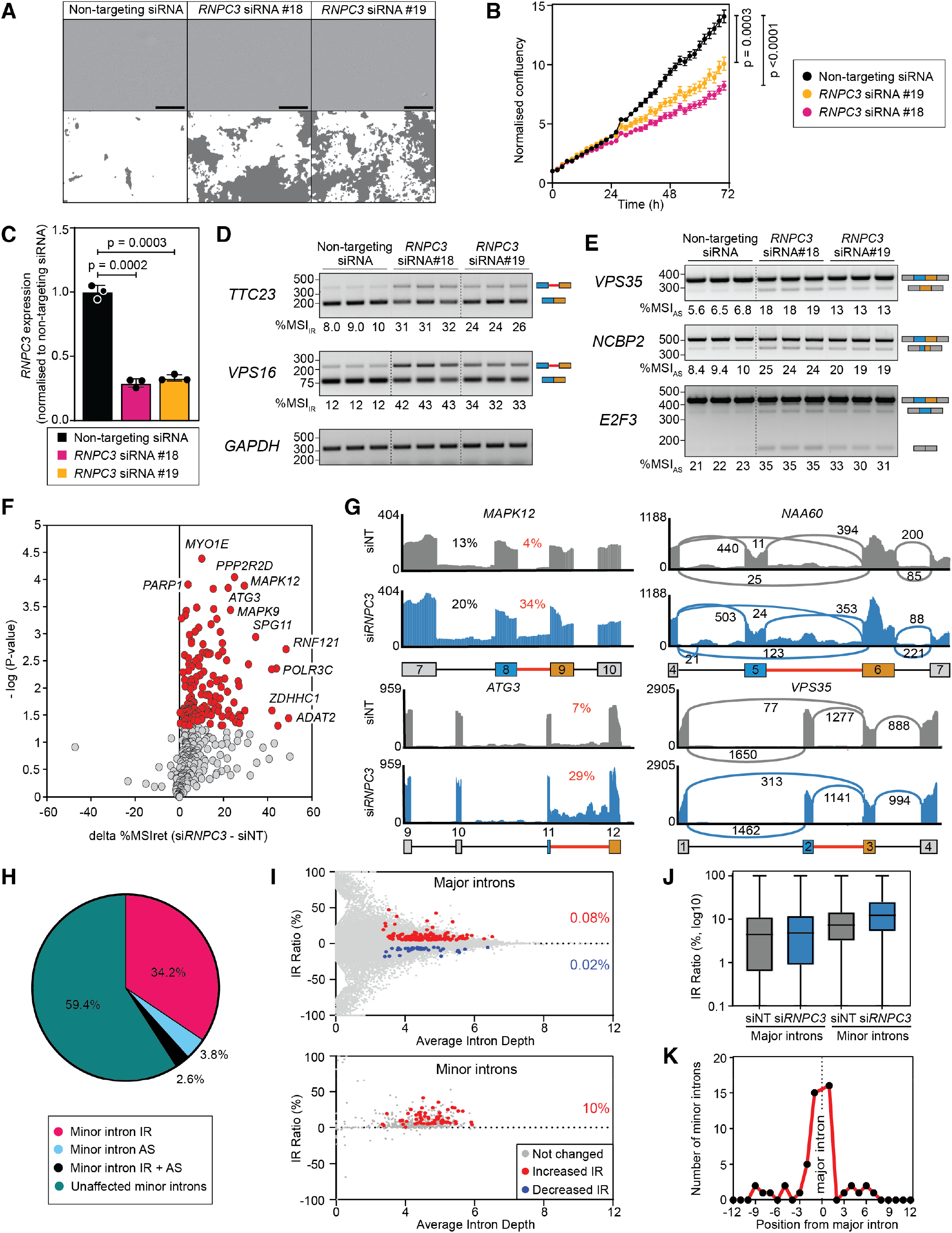
*RNPC3* knockdown impairs the growth of A549 cells and causes defects in the splicing of minor introns. **(A)** Representative images of A549 cells 72 h after siRNA transfection. Masking on phase objects using IncuCyte live cell image analysis shows that cells treated with si*RNPC3* for 72 h are less confluent than non-targeted (NT) siRNA treated cells. Scale bar is 400 μm. **(B)** Quantification of A549 cell growth following treatment with *RNPC3* and NT siRNAs for 72 h (25 images per well). Significance was assessed by one-way ANOVA (n=3). **(C)** RT-qPCR analysis shows marked knockdown of *RNPC3* transcripts in response to two independent *RNPC3* siRNAs. Data are represented by the mean ± SEM (n=3). **(D)** RT-PCR products of model minor introns showing intron retention and **(E)** alternative splicing in si*RNPC3*-treated A549 cells. *GAPDH* was used as loading control. Schematic depictions of the obtained amplicons are shown on the right with the minor intron in red and the upstream and downstream exons coloured blue and orange, respectively. % MSI_IR_ and %MSI_AS_ values were calculated using Image J. **(F)** Volcano plot of MIG transcripts exhibiting significant minor intron retention (solid red circles) in si*RNPC3*-treated cells. **(G)** Example RNAseq read coverage plots showing *RNPC3* knockdown increases minor intron retention in *MAPK12* and *ATG3* and produces alternative splicing in *NAA60* and *VPS35*. **(H)** Pie chart showing the percentage distribution of aberrant minor intron splicing events in si*RNPC3*-treated cells. **(I)** Transcriptome-wide IRFinder analysis showing median IR for all introns, irrespective of whether they are differentially retained or not (grey circles). Major introns (upper panel) showing significantly upregulated IR (solid red circles) and significantly downregulated IR (solid blue circles). Minor introns (lower panel) showing significantly upregulated IR (solid red circles). **(J)** Box plot of intron retention across all introns. **(K)** Retained major introns are found mostly in positions adjacent to minor introns.

### Knockdown of *RNPC3* expression disrupts minor intron splicing in human cancer cells

To determine the mechanisms underlying the reduced rates of A549 cell proliferation in response to *RNPC3* and *PDCD7* knockdown, we used RT-PCR analysis to identify disruptions in the splicing of minor introns (35,36). We found that siRNAs targeted to *RNPC3* caused aberrant splicing of some well-characterized minor introns, either through intron retention (IR) as seen for *TTC23* and *VPS16* (Figure 6D) or alternative splicing (AS) as seen for *VPS35, NCBP2* and *E2F3* (Figure 6E). These events were recapitulated in A549 cells treated with *PDCD7* siRNAs (Figure S5D) and HeLa cells treated with *RNPC3* and *PDCD7* siRNAs (Figure S5I).

To obtain a transcriptome-wide evaluation of gene expression and splicing changes that occur in A549 cells in response to 72 h of *Rnpc3* knockdown, we performed RNAseq. Using an intron retention pipeline designed to capture IR and AS of minor introns only (16,35), we found that of the 755 minor introns in the human genome, 51.9% showed evidence of retention while the remaining 48.1% did not. Of the retained minor introns, 59.4% showed no change in mis-splicing index (defined in Materials and Methods) in response to *RNPC3* knockdown (Figure 6F), 34.2% exhibited significantly elevated IR (*MAPK12* and *ATG3* in Figure 6G), 3.8% exhibited significantly increased AS (*NAA60* and *VPS35* in Figures 6G, S6D) and 2.6% exhibited elevated levels of both IR and AS (Figure 6H). We investigated IR in six MIGs by RT-qPCR and found that *MAPK8* expressed stable minor intron-containing transcripts not subjected to NMD, while *BRAF* expressed fewer transcripts overall, most likely because those containing minor introns were degraded by NMD (Figure S6E). Using gene ontology analysis, we identified molecular functions and biological processes associated with the incorrectly processed MIGs. ‘Cellular response to stress’ was the term most enriched with MIGs directly impacted by *RNPC3* knockdown (Figure S6F, Supplementary dataset 1).

We then performed a transcriptome-wide analysis of IR across all introns using IRFinder (13). Differential intron retention analysis identified 245 high-confidence introns that were differentially retained between si*RNPC3* and siNT transfected A549 cells (Figure 6I, Supplementary dataset 2). Of these, 185 were major introns and 60 were minor introns (Figure 6I). Of the 179,463 major introns detected, 148 (0.08%) exhibited increased IR in *RNPC3* knockdown cells and 0.02% exhibited reduced IR (Figure 6I upper panel). In contrast, of the 599 minor introns detected, 60 (10%) exhibited increased retention in *RNPC3* knockdown cells and no minor introns displayed less IR than in siNT-treated cells (Figure 6I lower panel). This equates to ∼125-fold enrichment of minor intron retention (10%) over major intron retention (0.08%) in *RNPC3* knockdown cells. The median retention level of all major introns (including the non-differentially retained introns) increased 1.05-fold upon *RNPC3* knockdown, while the median retention level of all minor introns increased almost 1.6-fold (Figure 6J), reiterating that the direct effect of *RNPC3* knockdown was primarily on minor introns. Of the retained major introns, one-third was found in transcripts containing one or more minor introns and almost two-thirds of these occupied positions flanking minor introns, independent of whether the minor intron was retained or not (Fig 6K). This phenomenon, which is likely to exacerbate the incorrect processing of the corresponding MIG transcripts, has been reported previously (16,35) and was also observed with our minor intron-specific IR pipeline (Figures 6G, S6G). In conclusion, depleting *RNPC3* in A549 cells compromises the integrity of minor intron splicing yet has a practically negligible impact on major introns except for some residing in MIGs.

### *RNPC3* knockdown causes transcriptome-wide differential gene expression in A549 cells

Of the 24,061 significantly expressed transcripts in our RNAseq dataset, 583 (2.4%) were MIGs, 23.7% (138) of which were significantly downregulated. In total, there were 1283 significantly downregulated DEGs and 985 significantly upregulated DEGs (Figure 7A, B). MIG abundance was significantly enriched in the downregulated set (10.8%), compared to the 2.4% expected if gene expression changes occurred randomly across the transcriptome (Figure 7B). Meanwhile, 3.2% of the upregulated DEGs were MIGs, not far from the expected 2.4% (Figure 7B). Approximately 90% of DEGs did not contain a minor intron, possibly signifying genes that were indirectly impacted by *RNPC3* knockdown consequent to a primary impact on MIGs.

**Figure 7.**
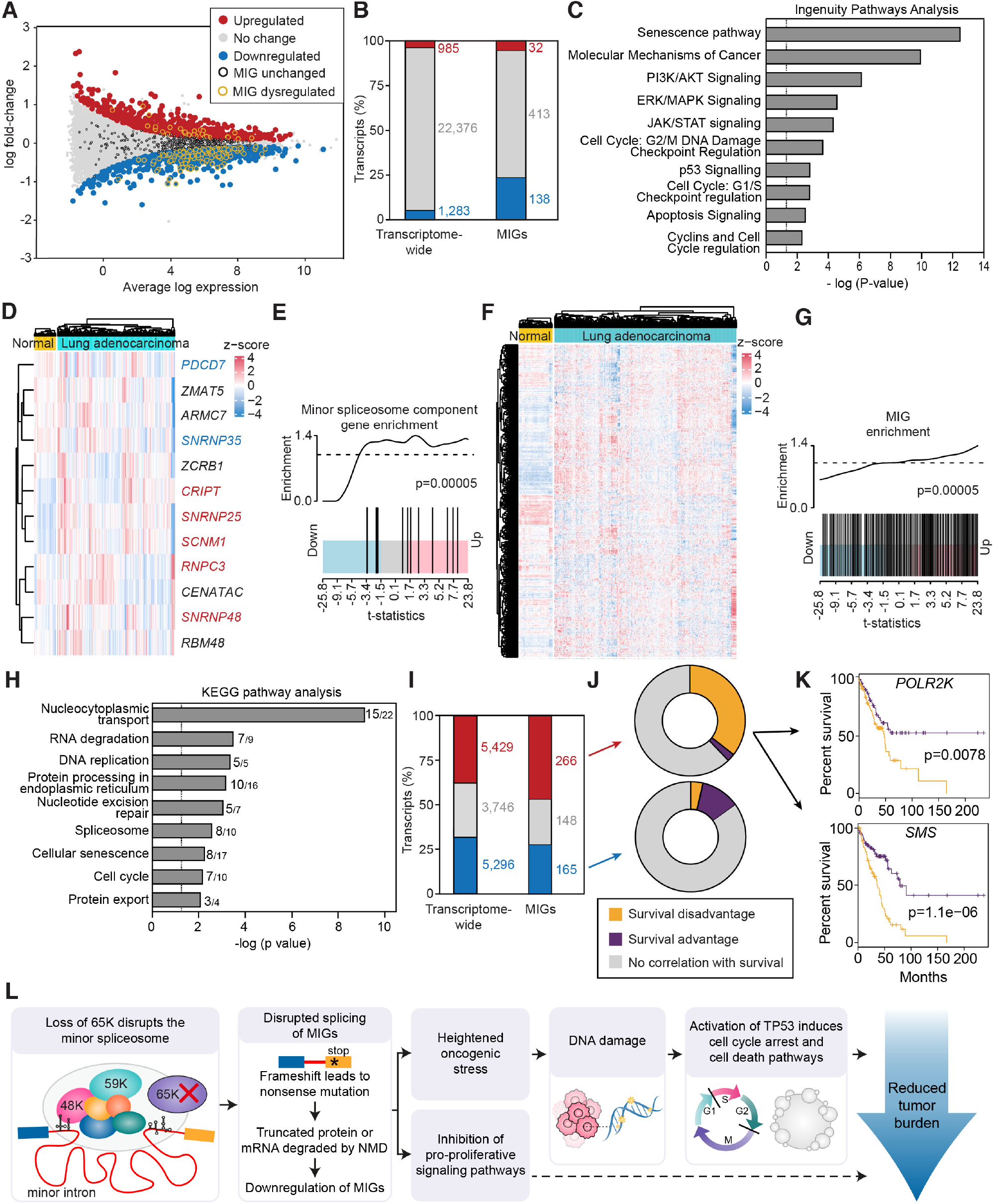
Knockdown of *RNPC3* results in differential expression of genes enriched in cancer related pathways. Transcripts encoding minor spliceosome-specific components and MIGs are upregulated in human lung cancer patients. **(A)** Scatter plot of differential gene expression between si*RNPC3* and siNT treated A549 cells. Of the 24,061 expressed transcripts, 2268 are DEGs (1286 downregulated shown in blue, 985 upregulated shown in red. The dysregulated MIGs are circled in yellow). **(B)** Differential gene expression analysis demonstrates a larger proportion of MIGs (10.8%; 138 out of 1283) are downregulated compared to the 2.4 % expected (583 expressed MIGs out of a total of 24,061 transcripts) if gene expression changes occurred randomly across the transcriptome (P<0.00001; Chi-square goodness of fit). **(C)** Selection of significantly enriched canonical pathways identified by IPA analysis of all affected genes (DEGs plus IR and AS affected MIGs). -log (*P*) values >1.3 (vertical line) signifies significant enrichment. **(D)** Heatmap of expression of minor spliceosome components in TCGA lung adenocarcinoma (LUAD) data. **(E)** Barcode plot of minor spliceosome components in LUAD samples compared to healthy tissue. **(F)** Heatmap displaying expression of MIGs in the TCGA LUAD dataset. **(G)** Barcode plot from gene set enrichment analysis of MIG expression compared to healthy tissue. All heatmaps show log-counts per million (logCPM) scaled for each gene. p-value for **E** and **G** = 5×10^−05^ as assessed by a roast gene set test with 9,999 rotations. **(H)** Upregulated MIGs are enriched in cancer relevant KEGG pathways. -log (*P*) values >1.3 (vertical line) signifies significant enrichment. The number of upregulated MIGs and total MIGs in each pathway are shown for each bar. **(I)** Differential gene expression analysis demonstrates almost half of expressed MIGs are upregulated in LUAD. **(J)** Survival analysis performed on the up and downregulated MIGs. **(K)** Example Kaplan-Meier survival plots of upregulated MIGs (*POLR2K* and *SMS*) associated with poor survival (n=120 patients/cohort). **(L)** Schematic diagram depicting the sequence of molecular and cellular events linking minor splicing disruption with reduced tumour burden.

Overall, ∼29% of MIG transcripts were differentially expressed by *RNPC3* knockdown (Figure S7A) and ∼71% were not. However, this 29% cohort of DE-MIGs did not include all the MIGs that were affected by IR and AS events (Figure S7B). Since these would preclude the production of functional proteins, we calculated that ∼40% of MIGs were affected by *RNPC3* knockdown, either through differential gene expression, intron retention and/or aberrant splicing. To identify the pathways directly impacted by this set of affected MIGs (DE-MIGs + IR MIGs + AS MIGs) we used Ingenuity Pathway Analysis (IPA). Of the top ten terms of ‘diseases and functions’, cancer was at the top (Figure S7C). We then used IPA to manually select canonical pathways significantly enriched with all affected genes (transcriptome wide DEGs + IR MIGs + AS MIGs) identifying terms relevant to this study such as cell cycle, senescence, DNA damage checkpoints, Tp53 signalling and the ERK/MAPK, PI3/AKT and JAK/STAT signalling pathways that drive cell proliferation and survival (Figure 7C; Supplementary dataset 1).

### Lung and liver cancers over-expressing minor splicing component genes and MIGs are associated with poor patient survival

To explore the clinical relevance of our findings, we interrogated the lung adenocarcinoma (LUAD) and liver hepatocellular carcinoma (LIHC) datasets of The Cancer Genome Atlas (TCGA) for gene expression patterns informative of minor splicing efficiency and patient survival. The LUAD dataset comprises 329 patient cancer transcriptomes and 59 healthy transcriptomes. Focusing first on the expression levels of the genes encoding the 12 unique minor splicing proteins, we found that 5 genes (red text), including *RNPC3*, were significantly upregulated in the LUAD dataset compared to their expression in normal lung tissue (Figure 7D, E). Turning to the expressed MIGs, we found that 45.9% were upregulated, equating to 4.9% of all upregulated transcripts in the LUAD dataset (Figure 7F, G). This 45.9% sub-set of upregulated MIGs was significantly enriched in KEGG pathway terms relevant to cancer and splicing such as nucleocytoplasmic transport, RNA degradation, DNA replication, spliceosome, cellular senescence and cell cycle (Figure 7H; Supplementary dataset 3).

We determined whether the expression of individual DE-MIGs was associated with patient outcome using Kaplan Meier survival analysis. We found that of the 266 MIGs exhibiting upregulated expression, 35% were associated with poor survival (orange in Figure 7J), including POLR2K and SMS (Figure 7K). Of the 165 downregulated MIGs, 85% exhibited no correlation with patient survival, 11.5% were associated with a survival advantage and 3.5% were associated with a poor survival outcome (Figure 7J). Analysis of the LIHC dataset, comprising 358 cancer transcriptomes and 50 healthy liver transcriptomes also revealed significantly upregulated expression of genes encoding minor spliceosome components and MIGs in LIHC tumours and that 65% of the upregulated MIGs were associated with poor survival (Figure S8).

## Discussion

Herein we show that the growth and spread of vigorously proliferating cancer cells are reliant on the integrity of minor splicing and that when this process goes awry, impaired splicing of MIGs leads to DNA damage and activation of a Tp53-dependent tumour suppressor-like transcriptional program that reduces tumour burden (Figure 7L). We found that transcripts encoding minor splicing components and MIGs are significantly upregulated in human lung and liver cancers and highly expressed MIGs are associated with poor patient survival, suggesting that cancer cells may employ mechanisms to increase the efficiency of minor splicing. We do not interpret this to mean that MIGs are cancer-causing oncogenes. Rather, we think that cancer cells expressing mutant KRAS proteins and other strong oncogenes require more efficient minor splicing and elevated expression of MIGs so that the molecular pathways that oncogenes depend on do not become rate-limiting.

Our results suggest that targeting the minor spliceosome may be useful for treating a spectrum of hitherto difficult-to-treat cancers. Inhibiting splicing in a cancer setting first attracted attention with the discovery that some cancer genomes carry point mutations in general splicing factor genes. For example, 20-30% of patients with myelodysplastic syndrome, a disease which frequently progresses to AML (16,37), harbour mutations in *SF3B1*. These mutations generally result in SF3B1 recognizing alternative splice sites leading to the production of aberrant proteins capable of promoting cell proliferation (38). Cancer cells bearing deleterious mutations in *SF3B1* are extremely sensitive to genetic or chemical disruption of splicing (39-41), and the development of pan-splicing inhibitors to kill these ‘sensitized’ cancer cells is a promising area of clinical research (42). However, this strategy may not be useful against cancers that do not carry sensitizing mutations, when higher doses of drug are likely to be required, causing unacceptable side-effects. We expect that inhibitors that are minor spliceosome-selective would leave a much smaller footprint of unwanted side effects.

Since the genes encoding BRAF, RAF1 and 11 out of 14 MAPKs contain minor introns, we thought that the pro-proliferative activity of the KRAS/RAF/MAPK pathway would rely heavily on efficient minor splicing, as indicated by the dashed arrow in Figure 7L. However, our results to date do not provide definitive evidence that mis-splicing of one or more BRAF/RAF1/MAPK transcripts is responsible for the reductions in tumour burden we observed in a variety of cancer settings. Instead, our data suggest that minor splicing is a cancer cell vulnerability that could be exploited to treat a broad spectrum of highly proliferative, oncogene-driven cancers by instigating changes in the expression of a more diverse set of cancer-relevant genes, such as those involved in DNA damage repair and replication. Such a mechanism may mean that targeting the minor spliceosome selectively would complement the use of targeted drugs that cancers often develop resistance to, leading to the successful treatment of a variety of cancers with a viable therapeutic index.

## Materials and Methods

All procedures performed on zebrafish and mice were conducted with the approval of the Animal Ethics Committees of the Walter and Eliza Hall Institute of Medical Research, The University of Melbourne and the Parkville Branch of the Ludwig Institute for Cancer Research, Australia. All mice were maintained on a C57BL/6 background. Zebrafish were maintained at 28°C on a 12 h light/12 h dark cycle according to standard husbandry procedures. Mice bearing constitutive and conditional mutant *Rnpc3* alleles were previously described (RRID:MGI:6276269) (11). Details of mice and zebrafish strains used in this study are available in Supplementary Methods.

### Inducing hepatocyte hyperplasia in zebrafish

To induce mutant Kras expression, *TO(kras*^*G12V*^) zebrafish larvae (21) were treated with 20 μg/ml doxycycline (Sigma, Cat#D9891) at 2 dpf in egg water with 0.003% 1-Phenyl-2-thiourea (PTU; Sigma, Cat#P7629) to suppress pigmentation. Egg water was changed at 5 dpf and fresh doxycycline (20 μg/ml) added. To quantitate liver volume, zebrafish larvae were anaesthetized with benzocaine (200 mg/L; Sigma, Cat#PHR1158) and mounted in 1% agarose. Image acquisition was performed using an Olympus FVMPE-RS multiphoton microscope with a 25x objective and Olympus FV30-SW software. Excitation wavelengths for GFP and dsRed were 840 nm and 1100 nm, respectively. Emission was detected at 550 nm and 580 nm, respectively. For volumetric analysis of whole livers, Z-stacks with step-size 2 μm, were imported into ImageJ (1.49v) or Imaris software.

### Cell death analysis in zebrafish

To assess apoptosis, 7dpf *TO(kras*^*G12V*^);*annexinV-mkate* zebrafish larvae were fixed in 4% paraformaldehyde (Thermo Fisher; Cat#28906) and the livers collected by microdissection. Image acquisition was performed using a Zeiss LSM 880 microscope with a 20x objective and ZEN software. Excitation wavelengths for mKate and GFP were 560 nm and 900 nm, respectively. Liver volume was quantified and 3D segmentation of the Annexin V-mKate signals was performed in FIJI.

### Cell cycle analysis in zebrafish

Live zebrafish larvae (7dpf) were incubated in 2mM EdU in egg water for 2 h followed by a further incubation in fresh egg water for 1 h. Larvae were euthanized using benzocaine (1000mg/L) prior to removal of the liver by dissection. EdU labeling was carried out using the Click-iT Edu Alexa Fluor 647 (AF647) imaging kit (Invitrogen, Cat#C10340) according to the manufacturer’s instructions. Livers were co-stained with Hoechst 33342 (1:250; Thermo Fisher, Cat#62249). Image acquisition was performed using an Olympus FVMPE-RS multiphoton microscope with excitation wavelengths of 950 nm and 1160 nm for Hoechst 33342 and AF647, respectively. The numbers of Hoechst 33342 and EdU positive cells were quantified using Arivis Vision4D software.

### AML cell lines

Phoenix cells (ATCC; RRID: CVCL_H717) transduced with MSCV-MLL-ENL-IRES-GFP (43) were cultured in Dulbecco’s modification of Eagle’s medium (DMEM; Gibco, Cat#11885084) supplemented with 10% fetal bovine serum (FBS; GE Healthcare Bio-Sciences, Cat#SH30088.03), 10% CO_2_ at 32°C for retrovirus production. Cells dissociated from E13.5 (embryonic day 13.5) mouse livers were cultured at 37°C with 10% CO_2_ in DMEM with 20% FBS, 100ng/ml murine stem cell factor (mSCF), 50ng/ml murine thrombopoietin (mTPO), 10ng/ml murine interleukin-6 (mIL-6) and 10ng/ml murine FMS-like tyrosine kinase 3 (mFlt-3), all produced in-house. Bone marrow derived secondary AML cells were cultured in DMEM with 10% FBS and mIL-3 (6ng/ml, Preprotech, Cat#213-13).

### *RNPC3* knockdown experiments in human cancer cells

Human lung adenocarcinoma, A549 cells (RRID: CVCL_0023) were cultured in RPMI medium supplemented with 1X GlutaMAX (Gibco, Cat#35050061) and 5% FBS. HeLa cervical cancer cells (RRID: CVCL_0030) were cultured in DMEM supplemented with 5% FBS. A549 cells were authenticated using small tandem repeat (STR) profiling (CellBank Australia, Report #19-338). HeLa cells were obtained directly from CellBank Australia and used for experimental work after 3 passages. A549 and HeLa cells were transfected with 50 nM of two independent human ON-TARGETplus siRNAs to RNPC3 (Dharmacon Inc., Cat#J-021646-18-0050 and J-021646-19-0050), PDCD7 (Dharmacon Inc., Cat#J-012096-05-0010 and J-012096-06-0010) or a non-targeting (NT) control siRNA (Dharmacon Inc., Cat#D-001810-01-05) using DharmaFECT 1 (Dharmacon Inc., Cat#T-2001-03) according to the manufacturer’s protocol. siRNA transfections were carried out in triplicate and treated as independent replicates thereafter. Cells were incubated immediately after transfection in an IncuCyte S3 live cell analysis system (Essenbioscience). A 5 × 5 imaging array was used to acquire 25 images per well every hour for 72 h for A549 cells and for 48 h for HeLa cells. Confluency over the time course was normalized to the first time point for each image acquired 1 h post-transfection.

### RNAseq of A549 cells

Total RNA was extracted from four independent transfections of A549 cells following 72 h treatment with si*RNPC3* or non-targeting (NT) control siRNA using TRIsure(tm) reagent (Bioline, Cat#38033). RNA integrity was assessed using a 2100 Bioanalyzer (Agilent). TruSeq Stranded mRNA Total Library Prep with Ribo-Zero Gold rRNA depletion (Illumina, Cat#20020598) was used to prepare cDNA libraries. Samples were sequenced on Illumina NovaSeq 6000, yielding >40M stranded paired-end 150bp reads per sample. RNAseq data are deposited at the GEO with Accession ID tba. See Supplementary Methods for differential gene expression analysis.

### Intron retention and alternative splicing analysis

To determine levels of intron retention (IR) and alternative splicing (AS) caused by *RNPC3* knockdown in A549 cells, RNAseq reads were aligned to the hg38 genome using Hisat2 with the – trim-5 1 option enabled. Minor intron retention and *de novo* alternative splicing events were then studied as described previously (4). Minor intron retention levels are reported as a mis-splicing index (MSI_ret_), which calculates the reads aligned to the 5’ and 3’ exon-intron boundaries as a ratio of the total number of reads spanning the canonical exon-exon junction. Other forms of alternative splicing (e.g. exon skipping) were quantified by calculating the ratio of spliced reads supporting an aberrant exon-exon junction over the total number of spliced reads aligning to the canonical exon-exon junction. Alternative splicing levels are reported as MSI_AS._ Significance of AS and IR events was assessed using a two-tailed T-test assuming unequal variance.

For analysis of global intron retention (both major and minor), we used IRFinder v1.2.0 following the default human genome pipeline (13). Differential IR (DIR) was called using an Audic and Claverie test. To ensure robust calling of DIR, only those significant DIR introns marked as “ clean” and with uniform or non-uniform coverage were considered. For an intron to be called as significantly retained in si*RNPC3* samples, we required the following: FDR ≤ 0.05, 100% coverage of the intron in si*RNPC3* samples, ≥ 10% IR ratio in siRNPC3 and ≥ 5% IR ratio over NT samples. Likewise, for an intron to be significantly retained in NT samples, we required FDR ≤ 0.05, 100% coverage of the intron in NT samples, ≥ 10% IR ratio in NT cells and ≥ 5% IR ratio over siRNPC3 samples. Minor introns were called based on the MIDB hg38 database (4).

### TCGA analysis

All count, sample and clinical data associated with the TCGA lung adenocarcinoma (LUAD) and liver hepatocellular carcinoma (LIHC) were downloaded using the GDC data portal. For both datasets, all genes with no symbol, duplicate gene entries, and gender specific genes were removed to avoid biases in the analysis. The data sets were then analyzed independently using the limma (44) and edgeR (45) software packages. See Supplementary Methods.

### Quantification and Statistical Analysis

Data are expressed as mean ± SEM unless indicated otherwise and the number of biological replicates indicating samples from individual animals for each experiment are stated in the figure legends. *P*-values were calculated using Student’s *t*-tests (two-tailed, followed by Welch’s correction) when comparing two groups, or tested by ANOVA followed by Tukey’s post-hoc test when comparing multiple groups. Survival data were plotted as Kaplan-Meier curves with significance calculated by Mantel-Cox log-ranked test. Chi-square goodness of fit test was used to evaluate whole transcriptome analysis and determine if MIG changes were statistically significant. All analysis was performed using GraphPad Prism V7.03 (GraphPad software) and P ≤ 0.05 was considered statistically significant.

## Acknowledgements

The authors thank Tyson Blanch, Cameron Mackey, Dora McPhee, Mark Greer, Lysandra Richards and Elizabeth Grgacic (zebrafish husbandry), Andrew Naughton, Melanie Asquith, Mel Pritchard, Faye Dobrowski, Emily Sutherland and Leanne Johnson (mouse husbandry), Qian Du, Julia Griesbach, Charlotte Burstroem and Samantha Eccles (technical assistance), Anne Voss and Tim Thomas (insightful discussions). This work was supported by the National Health and Medical Research Council of Australia (Project Grants 1024878 and 1161336 to JKH), Ludwig Cancer Research and a Victorian State Government Operational Infrastructure Support grant.

## Author contributions

### Conception and design

K. Doggett, S. Mieruszynski, B.B. Williams and J.K. Heath

### Development of methodology

K. Doggett, K.J. Morgan, S. Mieruszynski, B.B. Williams, A.M. Olthof, T.E. Hall, T.L. Putoczki, M. Ernst, K.D. Sutherland, Z. Gong and J.K. Heath

### Acquisition and analysis of data

K. Doggett, K.J. Morgan, S. Mieruszynski, B.B. Williams, A.M. Olthof, A.L. Garnham, M.J.G. Milevskiy, J. Coates, M. Buchert, R.J.J. O’Donoghue and J.K. Heath

### Writing, review, and/or revision of the manuscript

K. Doggett, K.J. Morgan, S. Mieruszynski, A.M. Olthof, A.L. Garnham, M.J.G. Milevskiy, R.N. Kanadia and J.K. Heath

### Study supervision

K. Doggett, B.B. Williams and J.K. Heath

### Funding Acquisition

J.K. Heath

