## Supplementary data for "The minor spliceosome offers a therapeutically viable target for the treatment of a broad spectrum of cancers"

**Supplementary Methods and References**

**Supplementary Figure 1, related to Figure 1**

**Supplementary Figure 2, related to Figure 2**

**Supplementary Figure 3, related to Figure 3**

**Supplementary Figure 4, related to Figure 4**

**Supplementary Figure 5, related to Figure 6**

**Supplementary Figure 6, related to Figure 6**

**Supplementary Figure 7, related to Figure 7**

**Supplementary Figure 8, related to Figure 7**

**Supplementary Table 1 – Primer sequences**

**Supplementary Dataset 1 - RNAseq analysis**

**Supplementary Dataset 2 - IRFinder**

**Supplementary Dataset 3 - TCGA KEGG analysis**

### Supplementary Methods

#### Mice

##### Lung adenocarcinoma model

8-16-week-old *Kras-LSL-G12D* mice (1) (Cat# JAX:008180) were exposed to adenoviral-Cre (AdCre) in the lung airway epithelia. AdCre:CaPi coprecipitates were prepared (2) and delivered intranasally to individual mice at a dose of 2.5 x $10^8$ plaque forming units in 50 µl MEM under isoflurane anesthesia. This treatment served to recombine both the *Kras-LSL-G12D* locus and *loxP*-flanked *Rnpc3* alleles when present.

To analyze the degree of hyperplasia in the lungs of $Kras^{LSLKrasG12D}$ mice, animals were anesthetized, their lungs inflated (250 mm-$H_2O$ pressure) and fixed by cardiac perfusion of phosphate-buffered 4% paraformaldehyde. Lungs were harvested, soaked overnight at $4^oC$ in the same fixative followed by embedding in paraffin. Histological sections (4 µm) were cut and stained with hematoxylin and eosin (H&E). Three left and right lung lobe sections with at least 100 µm between them were imaged using a Panoramic Scan II. Quantification of tissue area was performed using FIJI. Segmentation was carried out using Ilastik (Interactive learning and segmentation toolkit; https://www.ilastik.org). The area and number of tumor lesions from six slides/mouse were averaged to give one data point per mouse. Differences in lung tumor grade were based on criteria established by Nikitin and colleagues (3).

##### Gastric cancer model

$Gp130^{Y757F/Y757F}$ mice in which gastric adenoma development occurs spontaneously and with 100% penetrance by 100d (4,5) were used to investigate gastric cancer. We disrupted the *Rnpc3* locus in the glandular epithelium of the stomach by crossing $Rnpc3^{lox/lox}$;$Gp130^{Y757F/Y757F}$ mice with *Tff1-CreERT2* mice (6). We administered tamoxifen (Sigma, St. Loius, MO; 30mg/ml) to adult $Rnpc3^{lox/lox}$;$Gp130^{Y757F/Y757F}$;*Tff1-CreERT2* mice by oral gavage in two consecutive daily doses (150 µl). Stomachs were collected from mice euthanized at 100 d or 180 d of age, opened longitudinally, washed three times in PBS with vigorous shaking and pinned out on silicone-coated plates and photographed. Gastric adenomas were resected, weighed and either snap frozen for later molecular analysis or fixed in 10% buffered formalin solution (pH 7.4) overnight, prior to embedding in paraffin for immunohistochemical analysis.

**Generation and propagation of AML cells**

The MSCV MLL-ENL IRES GFP retroviral construct was obtained from Dr. Stefan Glaser (first described in (7). Transfection of Phoenix cells with this construct was achieved using 4 µl of Fugene (Promega, Cat#E2311) per µg of plasmid DNA and viral supernatants harvested 24 and 48 h later (8). Disaggregated fetal liver cells were virally transduced by spin infection with 4µg/ml polybrene (Sigma, Cat#H9268) on two consecutive days (9). $1 \times 10^6$ cells were transplanted into 6-8 weeks old sub-lethally γ-irradiated (7.5 Gy) C57BL/6 female mice, 24h prior to transplantation by tail-vein injection. Blood was collected from the retro-orbital plexus and cell counts were obtained with an Advia 2120 hematological analyzer to monitor disease onset. As each mouse reached the ethical endpoint and was euthanized, primary AML cells were harvested from the bone marrow and spleen and filtered through 100µm cell strainers to generate single cell suspensions. Red blood cell lysis was performed on spleen cell preparations. Primary AML cells were expanded by transplanting into secondary recipient mice (no prior irradiation). Secondary transplant cells were harvested from the spleen and bone marrow of at least three independent recipient mice per genotype and used to analyze the impact of disrupting the *Rnpc3* locus *in vitro* and *in vivo*. *UBC-CreERT2* (10) mediated recombination of the *Rnpc3* locus *in vitro* was achieved using treatment with 200 nM 4-OHT (Sigma, Cat#H7904). Cre-mediated recombination of the *Rnpc3* locus *in vivo* was achieved using 30 mg/ml TMX (Sigma, Cat#T5648) administered by oral gavage on day 13 and 14 following transplant. Kaplan Meier analysis was used to plot the survival of mice harboring the tertiary transplanted AML cells.

**Zebrafish lines**

Zebrafish harboring the *rnpc3* null allele were purchased from Znomics (RRID: ZFIN_ZDB-GENO-140806-3). In this mutant line, the *rnpc3* locus is disrupted by retroviral insertion into intron 1, resulting in undetectable mRNA transcripts (11). $tp53^{M214K/M214K}$ (also known as $tp53^{e7/e7}$ and referred to herein as $tp53^{m/m}$, has been previously described (RRID:ZFIN_ZDB-ALT-050428-2) (12). Tg(*fabp10:dsRed, ela3l:GFP*)$^{gz12}$, referred to herein as *2-CLiP* (RRID: ZFIN_ZDB-ALT-090424-3), expresses dsRed in the liver and GFP in the exocrine pancreas but carries no oncogenic transgenes or mutations (13). The Tg(*fabp10:rtTA2s-M2;TRE2:EGFP-kras*$^{G12V}$) line referred to as TO(*kras*$^{G12V}$) (RRID:ZFIN_ZDB-ALT-151022-1) (14) and the cell death reporter line, Tg(*actb2:SEC-Hsa.ANXA5-mKate2,cryaa:mCherry*)$^{uq24rp}$ was described previously (15). Foci of mKate fluorescence correspond to AnnexinV-mKate fusion proteins binding to exposed phosphatidylserine on the inner leaflets of plasma membranes in cells undergoing apoptosis.

**Genotyping**

Genomic DNA (gDNA) was extracted from zebrafish larvae by incubation at 95°C for 20 min in 50 µl of 50 mM sodium hydroxide (NaOH), followed by neutralization with 5 µl of 1 M Tris-HCl (pH8.0). gDNA for mouse genotyping and to assess efficiency of Cre-mediated deletion of exons 4 and 5 from the mouse *Rnpc3* locus, was extracted from tissues and cells of interest by lysis overnight at 55°C in DirectPCR buffer (Viagen Biotech, Cat#102-T) containing proteinase K (Viagen Biotech, Cat#503-PK). For PCR primer sequences see Supplementary Table 1.

**RNA analysis by RT-PCR and RT-qPCR**

Total RNA was extracted from independent pools of dissected zebrafish livers using the RNeasy Micro Kit (QIAGEN, Cat#74004). RNA integrity was assessed by a High Sensitivity RNA ScreenTape assay (Agilent, Cat#5067-5579) on a 2200 TapeStation. Total RNA was extracted from human and mouse cells and tissues using TRIsure™ reagent (Bioline, Cat#38033). RNA integrity was assessed using a 2100 Bioanalyzer (Agilent). cDNA was generated from 1-10 µg RNA using the Superscript III First Strand Synthesis System (Invitrogen, Cat#18080051) and oligo(dT) priming according to the manufacturer's instructions. RT-quantitative PCR (RT-qPCR) was performed using a SensiMix SYBR kit (Bioline, Cat#QT605-05) on an Applied Biosystems ViiA™7 Real-Time PCR machine. Zebrafish expression data were normalized by reference to *hrpt1, b2m* and *tbp*; mouse expression data were normalized by reference to *Gapdh* and *Hprt1* expression, and human to *GAPDH* and *CCND1* expression. LinRegPCR V11.0 was used for baseline correction, PCR efficiency calculation and transcript quantification analysis (16). To detect spliced transcripts by RT-PCR, primers were designed to amplify sequences spanning exon-exon borders. To detect minor intron retention by RT-qPCR, primers were designed to hybridize to sequences in an upstream (5') or downstream (3') exon and their adjacent minor intron (Primer sequences; Supplementary Table 1). Relative expression levels were calculated by the $2^{-\Delta\Delta Ct}$ method and all results were expressed as the mean ± SEM of at least three independent biological replicates.

**Western blot analysis and antibodies**

Samples of mouse and zebrafish tissues were lysed in RIPA buffer (20 mM Hepes, pH 7.9, 150 mM NaCl, 1 mM $MgCl_2$, 1% NP40, 10 mM NaF, 0.2 mM $Na_3VO_4$, 10 mM β-glycerol phosphate). Nuclear protein extracts were obtained from A549 cells following siRNA transfection using NE-PER Nuclear and Cytoplasmic Extraction Reagents (Thermo Fisher Scientific, Cat#78833) as per the manufacturer's protocol. All buffers were supplemented with cOmplete Proteinase and PhosSTOP inhibitors (Roche Cat#11836170001 & Cat#04906837001). Nuclear protein lysates were treated with 50 ng/µL DNase I (Worthington Biochemical, Cat#NC9199796), incubated for 30 min on ice and cleared by centrifugation at 13,000 rpm for 20 min at 4°C. The protein concentration of samples was determined by BCA protein assay (Thermo Fisher Scientific, Cat#23227). 25-50 µg of protein per lane were resolved on NuPAGE Novex Bis-Tris 4-12% polyacrylamide gels (Invitrogen, Cat#NP0321BOX). Mouse and human samples were transferred to Immobilon-FL PVDF membranes (Millipore, Cat#IPFL00010) and blocked in Odyssey Blocking Buffer (LI-COR, Cat#927 40000). Mouse gastric polyp blot was incubated with anti-phospho-p44/42 MAPK (ERK1/2) (Thr202/Tyr204, 1:1000; Cell Signaling Technology, Cat#4370) and anti-alpha tubulin (DM1A) (loading control) (1:2000; Cell Signaling Technology, Cat#3873). Human A549 blot was incubated with anti-U11/U12 snRNP 65K (1:500; Santa Cruz Cat#514951) and anti-TATA box binding protein (TBP, loading control) (1:2000; Abcam, Cat#ab28175). Membranes were incubated with secondary antibodies: IRDye 680LT donkey anti-rabbit (1:10000; LI-COR, Cat#925-68023) and IRDye 800CW donkey anti-mouse (1:10000; LI-COR, Cat#925-32212) and scanned using an Odyssey infrared imaging system (Licor). Zebrafish samples were transferred to nitrocellulose blotting membranes (Amersham, Cat#10600003), blocked with 5% BSA and incubated with primary antibodies anti-Tp53 (9.1), (1:500; Abcam, #ab77813) and anti GAPDH (14C10) (1:1000; Cell Signaling Technology, #2118). Secondary antibodies goat anti-mouse HRP (1:5000; DAKO, Cat#P0447) and goat anti-rabbit HRP (1:5000; DAKO, Cat#P0448) were incubated with membranes and signals developed using Amersham ECL Western Blotting Detection Kit (Cytiva, Cat#RPN2108) and imaged on a Chemidoc Touch (Biorad). Relative protein abundance was calculated based on normalized integrated intensity.

**Immunohistochemistry and antibodies**

Immunohistochemical analysis was carried out on unstained histological sections of mouse tissue using either anti-65K antibody (1:250; Abcam, Cat#ab90090) or anti-pERK (1:1000; Cell Signaling Technologies, Cat#4370). Prior to staining, antigen retrieval was performed on the sections by heating to 100$^{o}$C (microwave) in 10 mM pH6 sodium citrate buffer. Endogenous peroxidase activity was inhibited with 3% $H_2O_2$ (v/v; Thermo Fisher Scientific, Cat#H325-100). Non-specific antibody binding was blocked using 10% goat serum (w/v; Sigma, Cat#G9023). Primary antibody binding was performed overnight at 4°C in a humidified container. Sections were incubated with anti-rabbit biotinylated secondary antibody and Vectastain Elite ABC HRP reagent (Vector Laboratories, Cat#PK-

6100) for 30 min at RT. Signals were detected with the Liquid Diaminobenzidine substrate Chromogen System (Dako, Cat#K3468) prior to counterstaining with haematoxylin. All images were captured using an upright Nikon Eclipse 90i or a Panoramic Scan II.

**Cryosectioning and immunofluorescence microscopy analysis**

Dissected zebrafish livers were fixed in 4% PFA overnight at 4°C and washed with PBS/0.1% Tween 20 before incubation in 30% sucrose in PBS overnight at 4°C. Livers were aligned in a tissue mold, embedded in OCT and frozen on dry ice. Livers were sectioned at 10 µm intervals using a Thermofisher Scientific Microm HM550 cryostat. Sections were washed with PBS before blocking with 10% FCS in PBS/0.3% Triton X-100. Incubation with γ-H2AX (1:1000; gift of James Amatruda) antibody was performed at 4°C overnight, followed by 1 h at RT with anti-rabbit AF647 (1:500; Thermofisher Scientific, #A31573). Prolong Diamond Antifade Mountant with DAPI (Thermofisher #P36962) was used for slide mounting. A Zeiss LSM880 Fast Airyscan Confocal microscope with a 63x objective was used for image acquisition and image analysis was performed in ImageJ.

**Fluorescence activated cell sorting**

To determine the burden and phenotype of AML at the time of euthanasia, live peripheral blood and/or bone marrow cells were collected and stained with FluoroGold (Sigma, Cat#39286), anti-Gr-1 (RA6-8C5) and anti-Mac-1 (M1/70) antibodies (Walter and Eliza Hall Institute Monoclonal Antibody Facility). GFP fluorescence was used to detect AML cells expressing the MLL-ENL fusion protein. The viability of AML cells *in vitro* following treatment with 200 nM 4-OHT for timed intervals was assessed using Annexin V-Alexa Fluor 647 (Life Technologies, Cat#A23-204) and 4 µg/ml propidium iodide (Sigma, Cat#P4864) exclusion staining. Briefly, cells were washed once with balanced salt solution (150 mM NaCl, 3.7 mM KCl, 2.5 mM $CaCl_2$, 1.2 mM $MgSO_4$, 7.4 mM HEPES, NaOH, 1.2 mM $KH_2PO_4$ and 0.8 mM $K_2HPO_4$) containing 5% FBS and resuspended in the same medium containing the two reagents. Data was collected on an LSR-II flow cytometer (BD Biosciences) and cell viability analyzed using FlowJo v10.1 software (FlowJo LLC).

**Differential expression analysis of A549 cells by RNAseq**

All reads were aligned to the human genome, build hg38, using the Rsubread (17) software package and the align function (v2.4.3). In all cases at least 93% of all fragments (read pairs) mapped to the genome. All fragments overlapping genes were summarized into counts using Rsubread's featureCounts function. Genes were identified using Gencode annotation to the human genome (v37). Differential expression analyses between the si*RNPC3* and siNT conditions were then undertaken using the limma (v 3.46.0) (18) and edgeR (v 3.32.1) (19) software packages.

Prior to analysis, all genes labeled 'To Be Experimentally Confirmed' (TEC) were removed. Expression based filtering was then performed using edgeR's filterByExpr function with default parameters. A total of 24,644 genes remained. Sample composition was then normalized using the TMM method (20). To identify differentially expressed genes between the conditions, the data was first transformed to log-counts per million (logCPM) with associated precision weights using voom (21). Differential expression was then assessed using linear models and robust empirical bayes moderated t-statistics (limma-voom pipeline) (22). To increase precision, the linear models incorporated a correction for replicate. The false discovery rate (FDR) was controlled below 5% using the Benjamini and Hochberg method. The heatmap of the minor intron genes was generated using the pheatmap software package (v 1.0.12). Limma's removeBatchEffect function was first applied to the logCPM data prior to generating the heatmap to remove the replicate batch effect.

**Pathway analysis**

To identify biological functions that could be affected by aberrant minor intron splicing, all MIGs with significantly ($P<0.05$) elevated minor intron retention were submitted to g:Profiler. To identify pathways, diseases and functions affected upon si*RNPC3*, all affected genes (differentially expressed genes + MIGs with elevated retention and/or AS) were used for Ingenuity Pathway Analysis (IPA). Terms with $P<0.05$ were curated.

**Analysis of the LUAD dataset**

Prior to analysis, expression-based gene filtering was performed. All genes were required to achieve a CPM greater than 15 in at least 10 samples to be included, resulting in 14,471 genes being retained. Sample composition was then normalized using the TMM method. To distinguish differentially expressed genes between the cancer and normal tissue, the data was first transformed to logCPM. Differential expression was then assessed using linear models and robust empirical bayes moderated t-statistics with a trended prior variance (limma-trend pipeline) (21,22). The Benjamini and Hochberg method was applied to control the FDR below 5%.

**Analysis of the LIHC dataset**

Expression based gene filtering was first performed, requiring all genes to achieve a CPM greater than 2 in at least 10 samples. 17,650 genes were retained for downstream analysis. The TMM method was then applied to normalize sample composition. Differential expression between the cancer and normal samples was assessed using the limma-voom pipeline (21,22) and the FDR was controlled below 5% using the Benjamini and Hochberg method. Despite removal of the gender specific genes, a gender effect was observed in the data. The linear models therefore included an adjustment for gender to increase precision.

For both datasets, gene sets tests of the MIG genes were performed using ROAST (23) with 9,999 rotations. Barcode plots were drawn using limma's barcode plot function. Heatmaps were generated using the ComplexHeatmap (v 2.6.2) software package with row scaling first performed by pheatmap. Analyses of the KEGG database were performed using limma's kegga function. Overall survival analysis was performed through the GEPIA portal (http://gepia.cancer-pku.cn/). Patients were stratified based on the transcripts per million (TPM) of each gene-of-interest into low-expression (bottom 25% quartile) and high-expression cohorts (top 25% quartile) (21,22).

#### Supplementary Figures

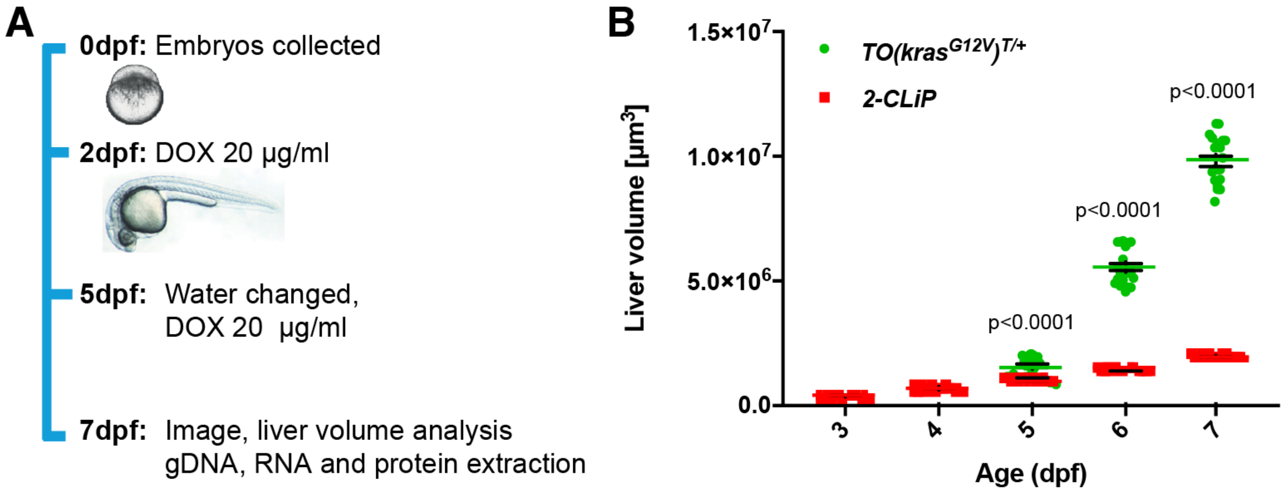


**Supplementary Figure 1.** Tissue-specific induction of *EGFP-kras*$^{G12V}$ expression in zebrafish hepatocytes causes an increase in liver volume. **(A)** Protocol for doxycycline (DOX) induction of mutant *EGFP-kras*$^{G12V}$ expression in developing zebrafish hepatocytes. DOX is added to the embryo medium at 2 and 5 d post-fertilization (dpf). Larvae are euthanized at 7 dpf and livers prepared for morphological and molecular analysis. **(B)** EGFP-Kras$^{G12V}$ protein expression in hepatocytes (green circles) is first detected by two-photon microscopy at 4 dpf and heralds a ~5-fold increase in liver volume over the next 3 d (5-7 days post-fertilization; dpf). Over the same period, normal liver growth, assessed with the *2-CLiP* transgenic line, which expresses dsRed constitutively in the liver, produces less than a 2-fold increase in liver volume. Data are expressed as mean ± SEM, n=20 per genotype. Significance was evaluated by multiple *t* tests using the Holm-Sidak method with $\alpha = 0.05$.

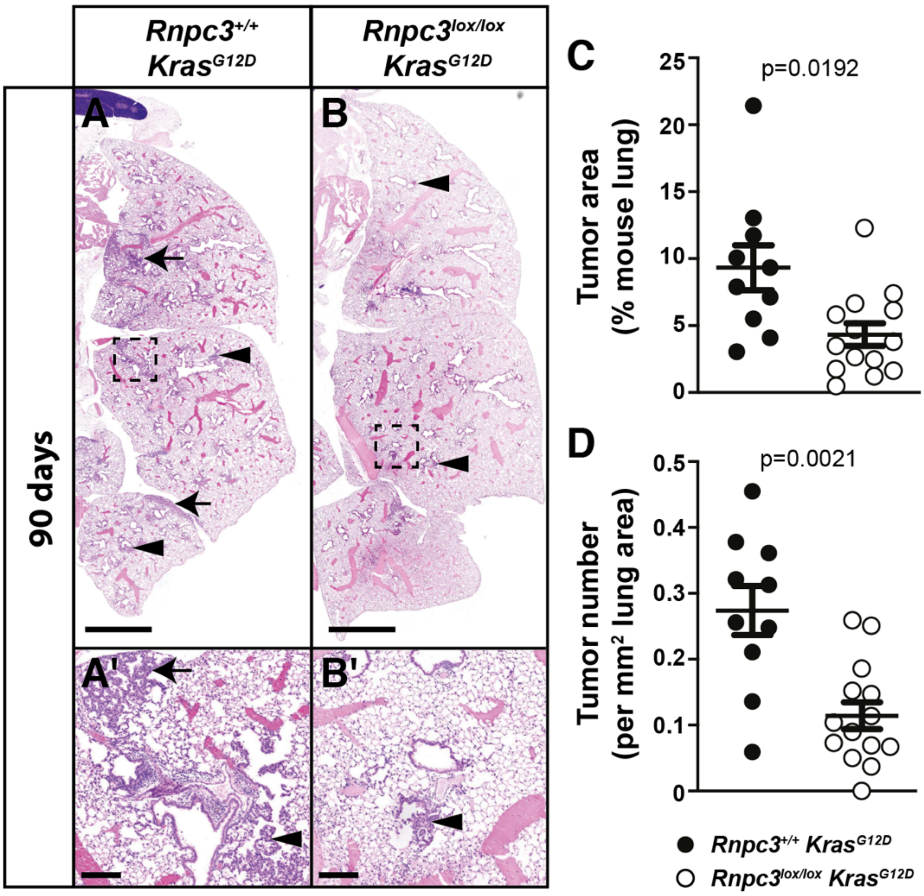


**Supplementary Figure 2.** Recombination of the *Rnpc3* locus reduces the area of atypical adenomatous hyperplasia (AAH) in a *Kras*$^{G12D}$-driven mouse model of lung adenocarcinoma. Adenoviral Cre recombinase was delivered to the airway epithelium of 8-11 weeks old *Rnpc3*$^{+/+}$;*Kras*$^{LSLKrasG12D}$ and *Rnpc3*$^{lox/lox}$;*Kras*$^{LSLKrasG12D}$ mice by intranasal injection. This treatment induced deletion of a *lox-stop-lox* cassette from the modified (knock-in) *Kras*$^{G12D}$ locus to initiate *Kras*$^{G12D}$ expression and recombination of *loxP*-flanked *Rnpc3* alleles. Histological analysis of lung hyperplasia was performed 90 d later. **(A)** Areas of AAH (arrows) and bronchial hyperplasia (arrowheads) were observed in the lungs of *Rnpc3*$^{+/+}$;*Kras*$^{LSLKrasG12D}$ and **(B)** *Rnpc3*$^{lox/lox}$;*Kras*$^{LSLKrasG12D}$ mice. **(A',B')** Higher magnification images of region in dashed box. **(C, D)** Quantitation of the frequency and area of the hyperplastic lesions revealed a significant decrease in AAH in *Rnpc3*$^{lox/lox}$;*Kras*$^{LSLKrasG12D}$ mice compared to *Rnpc3*$^{+/+}$;*Kras*$^{LSLKrasG12D}$ mice. Data are expressed as mean ± SEM, n=10 or 14 per genotype. Significance was assessed using an unpaired Student's *t* test with Welch's correction. Scale bar in a and b is 2 mm and in a' and b' is 200 µm.

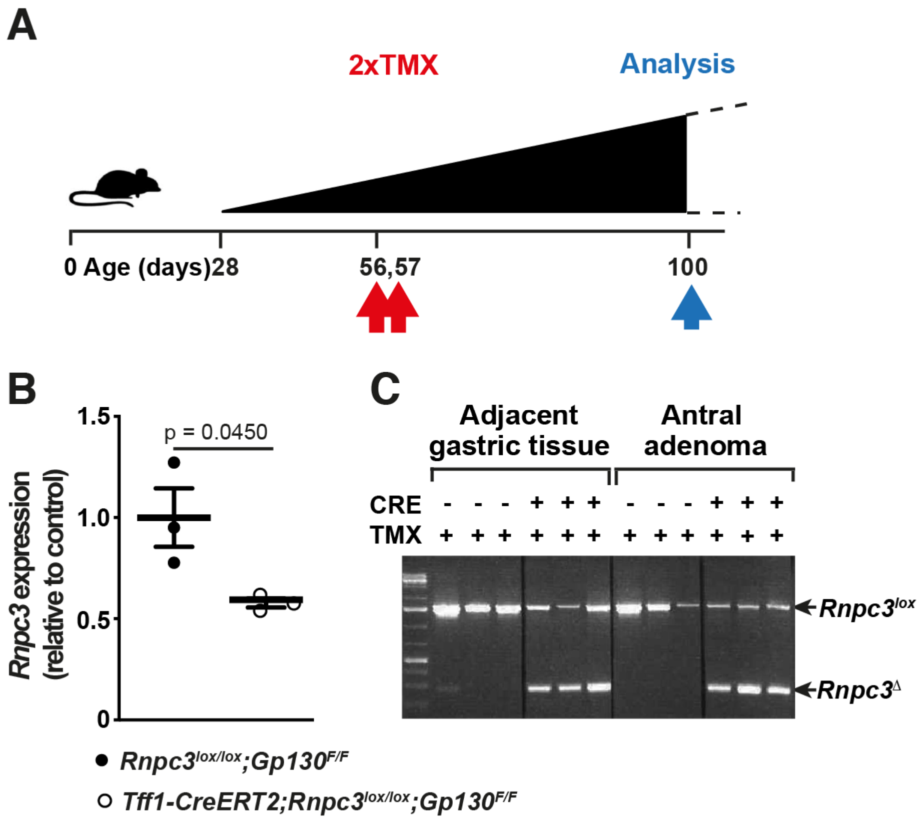


**Supplementary Figure 3.** Gastric epithelium-specific recombination of *Rnpc3*$^{lox}$ alleles in *Gp130*$^{F/F}$ mice. **(A)** Schematic diagram indicates that gastric adenoma formation in *Gp130*$^{F/F}$ mice is first evident from ~28d of age. Treatment with tamoxifen (2 x TMX) on days 56 and 57 (red arrows) is designed to induce recombination of *Rnpc3*$^{lox}$ alleles in developing gastric adenomas. The experiment is terminated at 100d when adenomas are weighed and subjected to molecular analysis (blue arrow). **(B)** RT-qPCR analysis reveals approximately 50% less *Rnpc3* mRNA in single antral adenomas from 3 independent TMX-treated *Rnpc3*$^{lox/lox}$*;Gp130*$^{F/F}$ mice harbouring the *Tff1-CreERT2* transgene (white circles)**,** compared to antral adenomas from control *Rnpc3*$^{lox/lox}$*;Gp130*$^{F/F}$ mice (black circles, mean set at 1). Significance was evaluated with a two-tailed Student's *t* test. **(C)** PCR of genomic DNA demonstrates the presence of both recombined (*Rnpc3*$^{\Delta}$) alleles (lower band) and unrecombined *Rnpc3*$^{lox}$ alleles (upper band) in normal glandular stomach and adenomas harvested from TMX-treated, *Tff1-CreERT2;Rnpc3*$^{lox/lox}$*;Gp130*$^{F/F}$ mice, n=3.

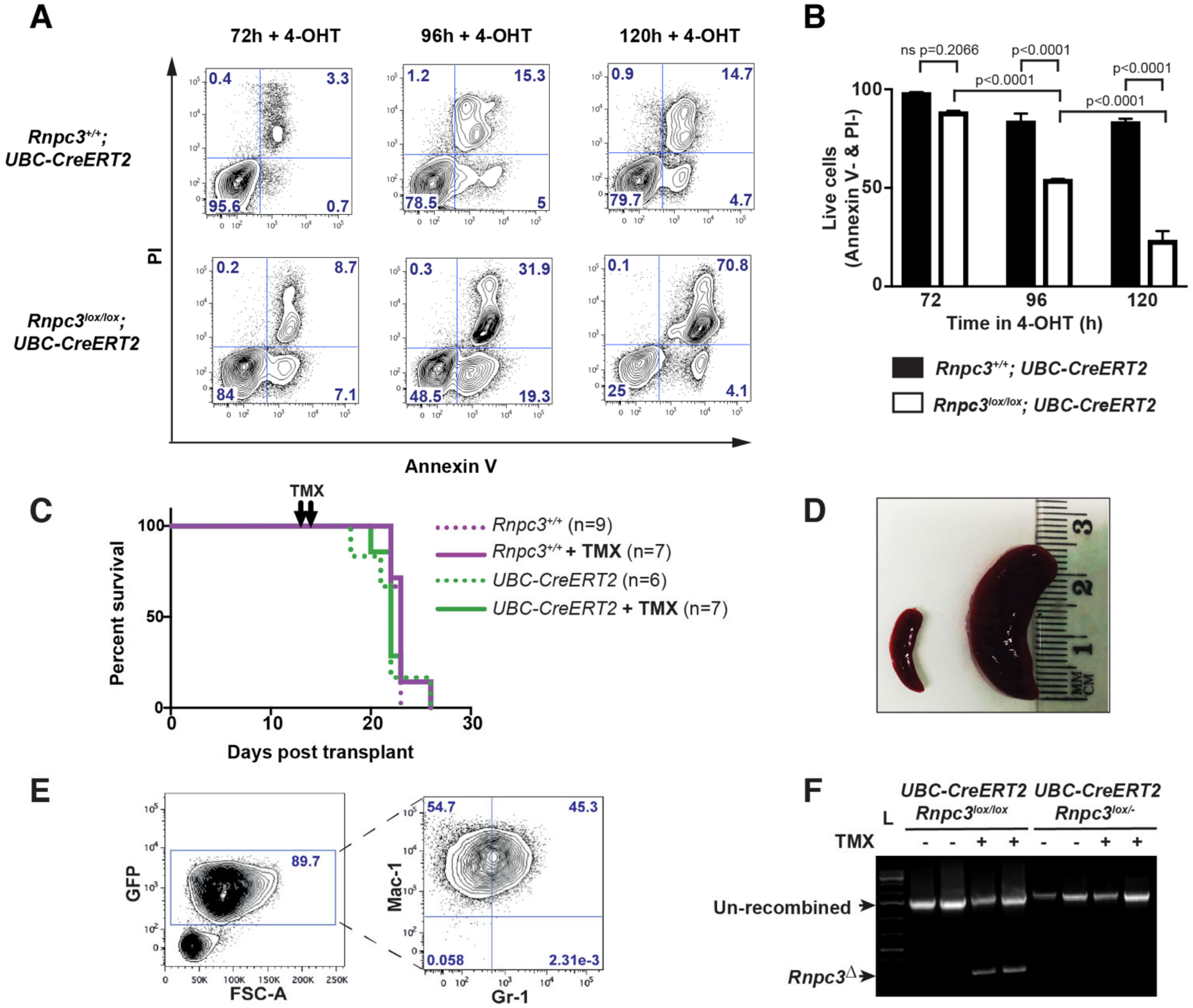


**Supplementary Figure 4.** Recombination of the *Rnpc3* locus in MLL-ENL AML cells *in vitro* reduces cell viability. Secondary transplants of MLL-ENL AML cells containing the *UBC-CreERT2* transgene in the presence (*Rnpc3$^{lox/lox}$*) and absence (*Rnpc3$^{+/+}$*) of *loxP*-flanked *Rnpc3* alleles were cultured for 72, 96 and 120 h in presence of 4-hydroxytamoxifen (4-OHT). **(A)** Representative fluorescence-activated cell sorting (FACS) plots of cell viability using AnnexinV and PI staining. Numbers in each quadrant indicate the percentage of cells in each category. A time-dependent decrease in viable *Rnpc3$^{lox/lox}$* cells (AnnexinV$^-$ and PI$^-$) is evident following 4-OHT treatment, such that by 120h only 25% of live cells remain, compared to 80% of *Rnpc3$^{+/+}$* cells also treated with 4-OHT. **(B)** Quantitation of the percentage of viable (AnnexinV$^-$, PI$^-$) cells over time enumerated by FACS. Percentage viable cells expressed as mean $\pm$ SEM, n=3 for all cohorts. Significant differences were assessed with a one-way ANOVA with Tukey's multiple comparison test. **(C)** Kaplan Meier plot depicting the survival of mice harbouring tertiary transplants of MLL-ENL AML cells with control genotypes (i.e. no *Rnpc3*$^{lox}$ alleles) in the presence (solid line) and absence (dotted line) of TMX treatment 13 and 14d (black arrows) following transplantation. There is no significant difference in median survival (22-23d) between all cohorts, n=6-9. Significance was assessed with a Mantel-Cox test. **(D)** At the ethical endpoint of the experiment, all recipient mice displayed splenomegaly independent of AML cell genotype. Image depicts a normal spleen from a WT mouse not transplanted with AML cells next to a spleen harvested from a mouse transplanted with *UBC-CreERT2*;*Rnpc3*$^{lox/lox}$ AML cells and treated with TMX. **(E)** FACS analysis of cells harvested from the bone marrow of a TMX-treated mouse transplanted with *UBC-CreERT2*;*Rnpc3*$^{lox/lox}$ AML cells. The GFP$^{+}$ cell population comprises cells that are both Mac-1$^{+}$ and Gr-1$^{+}$ positive, consistent with an immature myeloid phenotype. **(F)** Genomic analysis of the *Rnpc3* locus in tertiary transplanted *UBC-CreERT2*;*Rnpc3*$^{lox/lox}$ AML cells collected from TMX-treated mice at the ethical endpoint of the experiment reveals the presence of unrecombined and recombined *Rnpc3* alleles. Meanwhile *UBC-CreERT2*;*Rnpc3*$^{lox/-}$ AML cells collected from TMX-treated mice contain no *Rnpc3*$^{\Delta}$ alleles, indicating that *Rnpc3*$^{\Delta/-}$ cells did not survive the experiment.

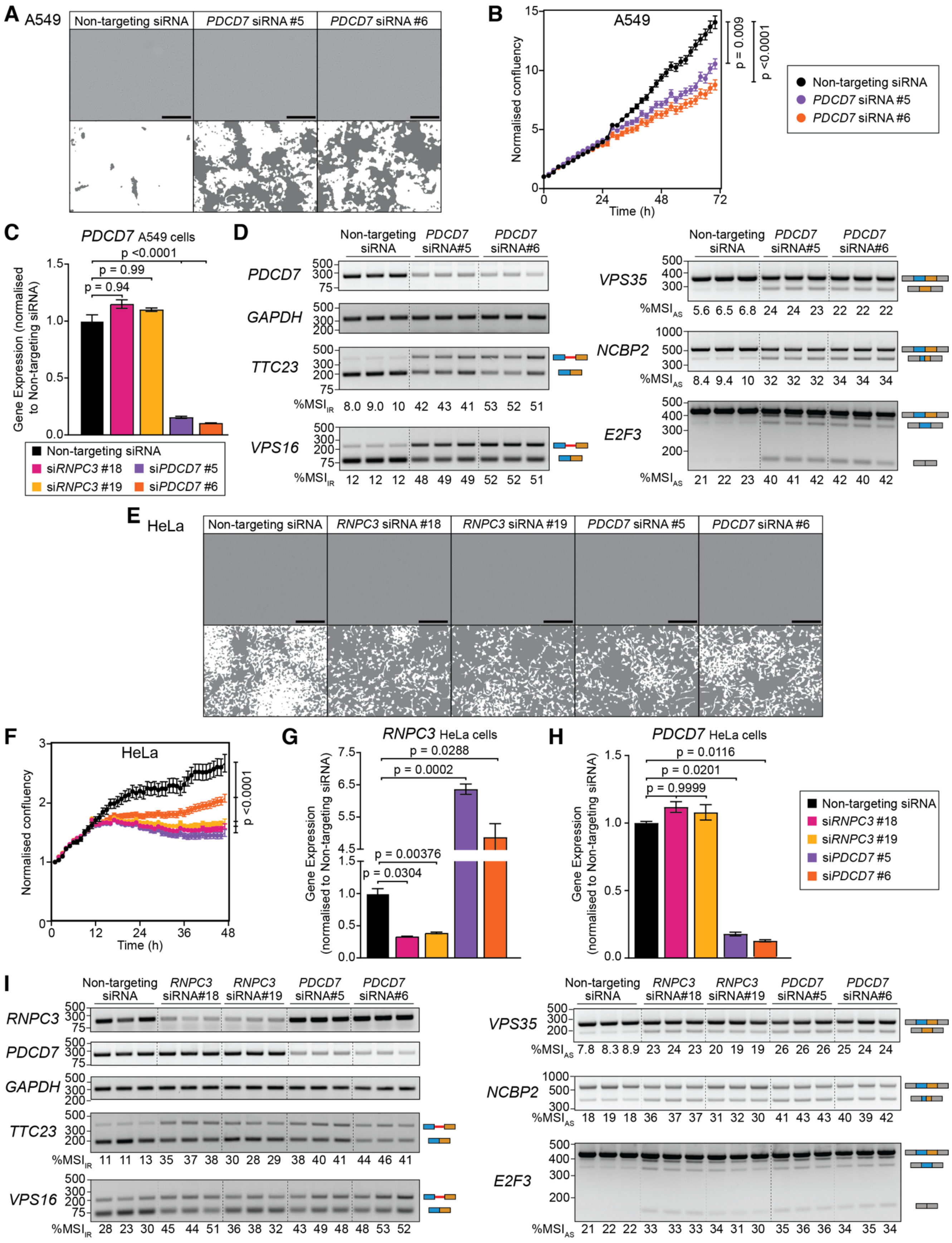


**Supplementary Figure 5.** siRNA knockdown of minor spliceosome components in human cancer cell lines restricts cell growth and induces MIG splicing changes. **(A)** A549 cell images 72 h after transfection with non-targeting or *PDCD7* siRNAs and masking for cell growth quantification. Grey areas correspond to absence of cells. **(B)** Reduced proliferation of *PDCD7* siRNA-treated A549 cells quantified from confluency curves generated in an IncuCyte live cell analysis system over 72 h. **(C)** RT-qPCR showing specificity of two independent siRNAs targeted to *PDCD7* in A549 cells for 72 h. **(D)** RT-PCR of A549 cells treated with *PDCD7* siRNAs showing *PDCD7* knockdown and MIG splicing changes. Schematics depicting splicing products are shown on the right with the minor intron in red and the upstream and downstream exons coloured blue and brown, respectively. % $MSI_{IR}$ and $\%MSI_{AS}$ calculated using Image J. **(E)** HeLa cell images 48 h after non-targeting, *RNPC3* or *PDCD7* siRNA treatment. **(F)** Both *RNPC3* and *PDCD7* siRNAs significantly restrict HeLa cell proliferation over 48 h. **(G)** RT-qPCR of HeLa cells upon *RNPC3* and **(H)** *PDCD7* knockdown. **(I)** RT-PCR analysis of MIG splicing changes in HeLa cells. Scale bar in all images is 400 µM. RT-PCR and RT-qPCR data are represented as mean ± SEM (n=3). Significance was assessed by one-way ANOVA with Tukey's multiple comparison test.

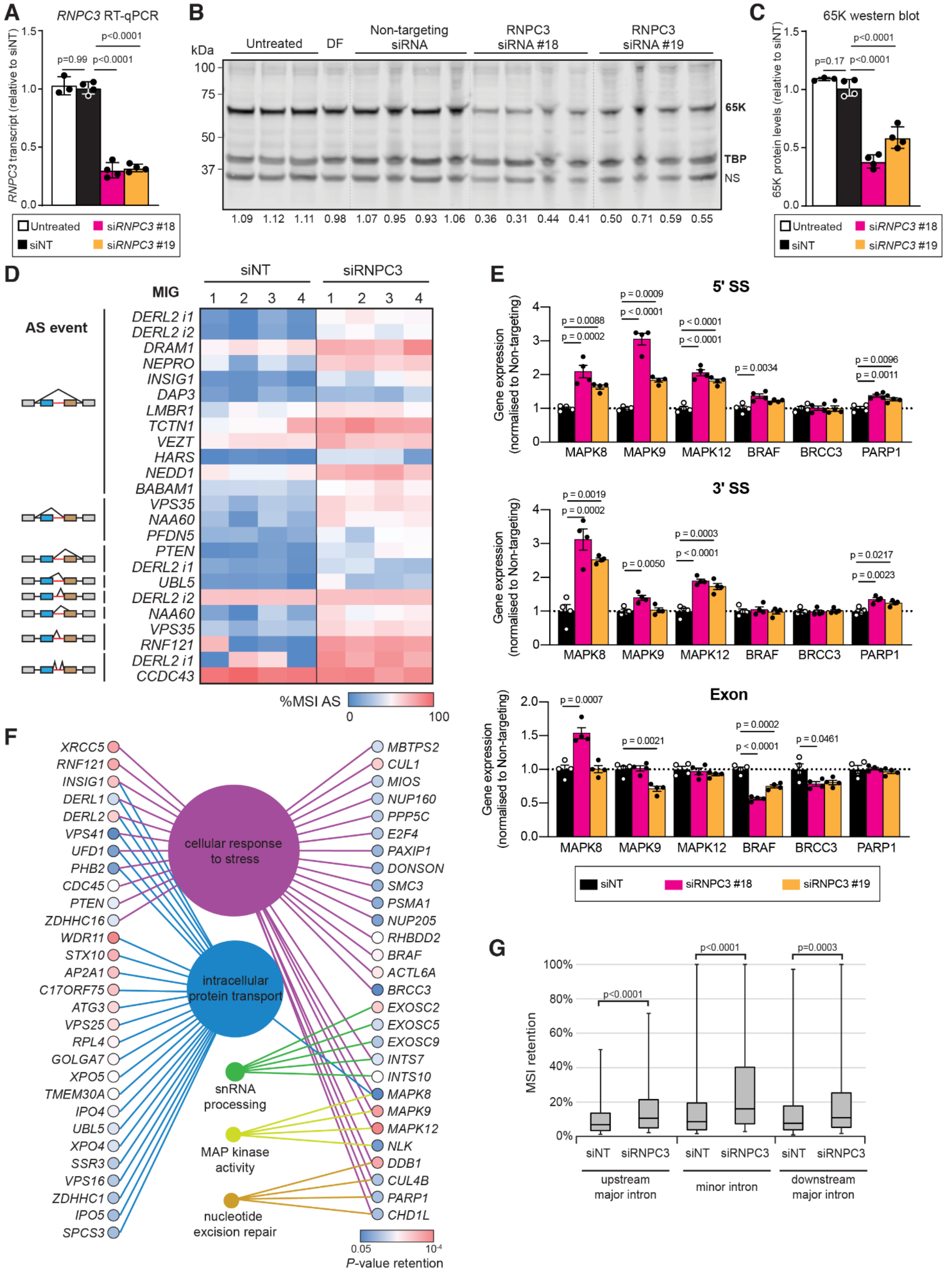


**Supplementary Figure 6.** RNAseq analysis of A549 cells treated with si*RNPC3* for 72 h reveals aberrant splicing of minor intron-containing genes (MIGs). **(A)** RT-qPCR analysis of *RNPC3* transcripts in A549 cells treated with si*RNPC3* #18 and #19 for 72 h demonstrates >70% knockdown. **(B)** Corresponding Western blot analysis of nuclear protein lysates from the same A549 cells shows 65K and TBP proteins. **(C)** Quantification of blot shown in **(B)** reveals a 62% and 45% reduction in 65K protein in response to si*RNPC3* #18 and si*RNPC3* #19, respectively. DF = DharmaFect (transfection reagent alone), NS = Non-specific band. Values shown are normalized by reference to the TBP loading control and relative to the non-targeting control. **(D)** Heatmap of identified MIG alternative splicing (AS) events. The schematized AS events are represented on the left with the minor intron in red and the flanking upstream and downstream exons in blue and brown, respectively. **(E)** RT-qPCR evaluation of a selection of MIG IR events across the minor intron 5'ss (splice site), 3'ss or of total transcript expression (exon primers spanning a major intron) upon transfection with two independent siRNAs to *RNPC3*. **(F)** Selected enriched GO terms of MIGs with significant intron retention. **(G)** Box plot of intron retention at minor introns and their upstream and downstream major introns. Significance was assessed by Mann-Whitney U test. The data in **(A)**, **(C)** and **(E)** are represented as mean ± SEM (n=4). Significance was assessed by one-way ANOVA with Tukey's multiple comparison test, $p<0.05$ was considered significant.

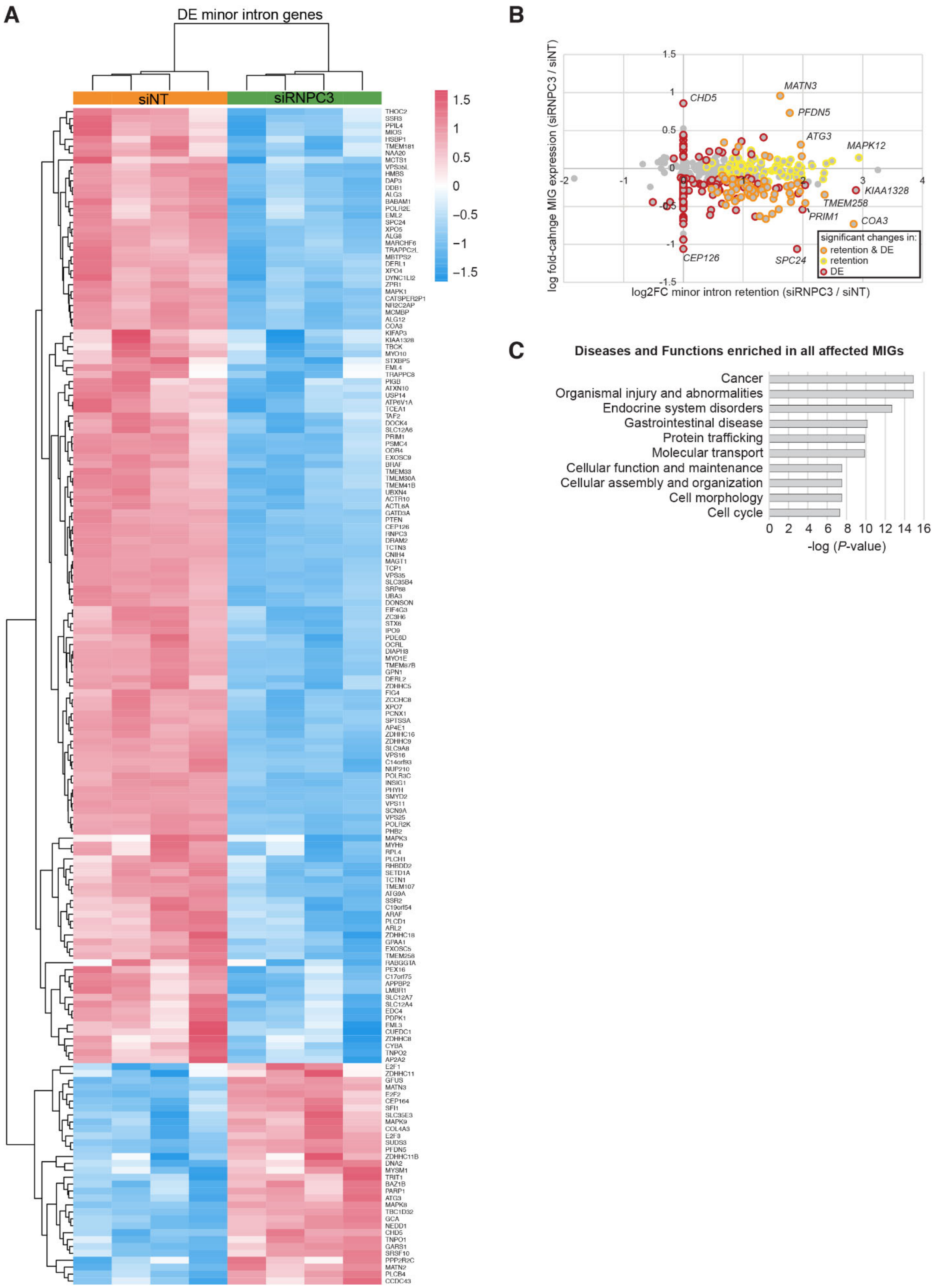


**Supplementary Figure 7.** A549 cells treated for 72 h with *RNPC3* siRNA exhibit differential expression of genes and MIGs that are enriched in cancer. **(A)** Heatmap of all identified differentially expressed MIGs, showing logCPM corrected for replicate and scaled for each gene. **(B)** Log-fold change plot of all MIGs highlighting those that exhibit significant changes in intron retention, differential expression or both when *RNPC3* is knocked down. **(C)** Top 10 enriched diseases and functions identified by IPA analysis of all affected MIGs.

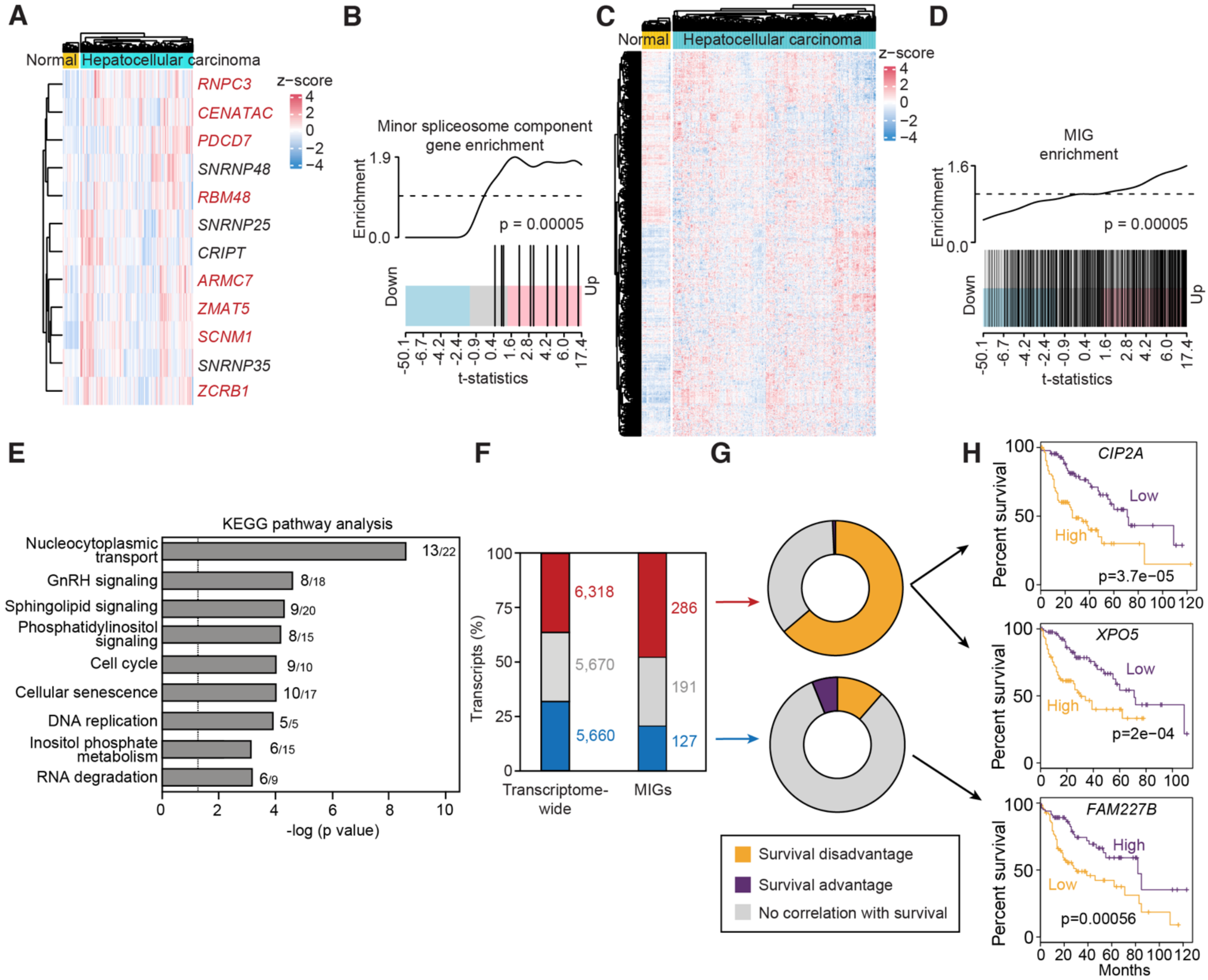


**Supplementary Figure 8.** Transcripts encoding minor spliceosome-specific components and MIGs are upregulated in the TCGA liver hepatocellular carcinoma (LIHC) dataset. **(A)** Heatmap of expression of minor spliceosome components. **(B)** Barcode plot of minor spliceosome components. **(C)** Heatmap displaying expression of MIGs. **(D)** Barcode plot from gene set enrichment analysis of MIG expression compared to healthy tissue. **(E)** Upregulated MIGs are enriched in cancer relevant KEGG pathways. -log (*P*) values >1.3 (vertical line) signifies significant enrichment. The number of upregulated MIGs and total MIGs for each pathway are denoted besides each bar. **(F)** Differential gene expression analysis demonstrates almost half of expressed MIGs are upregulated in LIHC. **(G)** Survival analysis performed on the up and downregulated MIGs. **(H)** Example Kaplan-Meier analysis of upregulated MIGs (*CIP2A* and *XPO5*) associated with survival disadvantage and downregulated MIG (*FAM227B*) associated with survival advantage (n = 1200 patients/cohort). All heatmaps log-counts per million (logCPM) scaled for each gene. P-value for e and g = $5x10^{-05}$ as assessed by a roast gene set test with 9,999 rotations.
